# SABER enables highly multiplexed and amplified detection of DNA and RNA in cells and tissues

**DOI:** 10.1101/401810

**Authors:** Jocelyn Y. Kishi, Brian J. Beliveau, Sylvain W. Lapan, Emma R. West, Allen Zhu, Hiroshi M. Sasaki, Sinem K. Saka, Yu Wang, Constance L. Cepko, Peng Yin

## Abstract

Fluorescent *in situ* hybridization (FISH) reveals the abundance and positioning of nucleic acid sequences in fixed samples and can be combined with cell segmentation to produce a powerful single cell gene expression assay. However, it remains difficult to label more than a few targets and to visualize nucleic acids in environments such as thick tissue samples using conventional FISH technologies. Recently, methods have been developed for multiplexed amplification of FISH signals, yet it remains challenging to achieve high levels of simultaneous multiplexing combined with high sampling efficiency and simple workflows. Here, we introduce signal amplification by exchange reaction (SABER), which endows oligo-based FISH probes with long, single-stranded DNA concatemers that serve as targets for sensitive fluorescent detection. We establish that SABER effectively amplifies the signal of probes targeting nucleic acids in fixed cells and tissues, can be deployed against at least 17 targets simultaneously, and detects mRNAs with high efficiency. As a demonstration of the utility of SABER in assays involving genetic manipulations, we apply multiplexed FISH of reporters and cell type markers to the identification of enhancers with cell type-specific activity in the mouse retina. SABER represents a simple and versatile molecular toolkit to allow rapid and cost effective multiplexed imaging.

## INTRODUCTION

Fluorescent *in situ* hybridization (FISH) allows researchers to interrogate the subcellular distribution of RNA and DNA molecules in fixed cells and tissues. First introduced in the late 1960s, *in situ* hybridization methods^1^ and their fluorescent variants^2–5^ utilize detectable nucleic acid probes that can be programmed to hybridize to complementary cellular targets. FISH assays are integral to many fields and are used for diverse applications such as the diagnostic detection of chromosomal abnormalities,^6^ the interrogation of three-dimensional genome orga-nization,^7, 7^ and the quantitative analysis of gene expression.^9, 9^

Despite their widespread use, conventional FISH approaches are still limited by technical considerations. For instance, it can be challenging to visualize hybridization events in complicated environments such as tissue samples due to inefficiencies in probe penetration and light collection. Accordingly, several approaches have been developed to amplify the intensity of quantitative FISH signals. These strategies include the targeted deposition of detectable reactive molecules around the site of probe hybridization,^11–14^ the targeted assembly of ‘branched’ structures composed of DNA^15–17^ or locked nucleic acid (LNA) molecules,^18^ the programmed *in situ* growth of detectable concatemers by enzymatic rolling circle amplification (RCA)^19, 19^ or hybridization chain reaction (HCR),^21–25^ and the assembly of topologically catenated DNA structures using serial rounds of chemical ligation.^26^

Using conventional FISH approaches, it is also difficult to examine more than a few targets in the same sample due to the relatively small number of spectrally distinct imaging channels available on fluorescent microscopes. In order to overcome this limitation, methods have been introduced that rely on creating additional ‘colors’ by labeling the same target with multiple fluorophores in defined combinations^27, 27^ and ratios,^29^ allowing the unambiguous detection of all 24 human chromosomes on metaphase spreads^27–29^ and in interphase nuclei^7^ and >30 mRNA targets in budding yeast.^30^ Additional approaches have been introduced that utilize serial rounds of imaging, label removal, and re-labeling of distinct targets, enabling researchers to image potentially unlimited numbers of targets. These strategies have been used to visualize up to 40 distinct chromosomal regions in tissue culture cells by imaging each target region just once with a single fluorophore.^31^ Multi-round combinatorial labeling strategies, which allow an exponential number of unique, non-overlapping targets to be visualized in a linear number of rounds, have been used to image 12 mRNA targets in budding yeast,^32^ >1000 mRNA targets in tissue culture cells,^33^ and intronic sequences^34^ of >10,000 nascent RNAs in tissue culture cells.^35^ However, because these strategies require spatially separated targets, only much more modest multiplexing levels have been demonstrated with arbitrarily dense targets.^36^ In principle, all of these multiplexing strategies could benefit from signal amplification, which could reduce requirements on expensive microscopy setups, shorten imaging times, and potentially reduce cost by lowering the number of probes required. This amplification would be particularly relevant in the context of thick tissues, which can suffer from high levels of autofluorescence, light scattering, and optical aberration that can make signal detection challenging.

Excitingly, recent studies have begun to integrate serially multiplexed imaging with signal amplification to detect targets in tissue samples. One approach, seqFISH,^32^ combined HCR with a combinatorial multi-round labeling strategy to visualize up to 249 mRNA targets in 15µm mouse brain tissue sections.^37^ However, design of orthogonal HCR concatemers is difficult due to the challenge of designing multiple non-interfering programmed reaction pathways to operate *in situ*, and only 5 orthogonal HCR concatemers have been demonstrated. As a result, only 5 barcodes could be deployed simultaneously in each sequential round, with each round consisting of concatemer digestion (4 hours), probe hybridization (overnight), amplification (∼1 hour), and imaging.^37^ Another approach, STARmap, used a novel form of RCA coupled with *in situ* RNA sequencing^38^ to detect up to 28 mRNA targets simultaneously in cleared 150µm mouse tissue sections and up to 1,020 spatially separated targets in thin (single cell layer) sections.^39^ While RCA-based methods in principle overcome the limitation of HCR by enabling highly multiplexed simultaneous amplification, detection efficiencies for RNA transcripts remain comparable to single-cell RNA sequencing (6-40%) in the best case (STARmap) and are even lower for other RCA-based methods such as FISSEQ (0.01-0.2%) and padlock probe-based designs (5-30%),^39^ perhaps due to the complexity of controlling parallel enzymatic reactions *in situ*. Furthermore, strategies combining signal amplification and high levels of multiplexing remain challenging for many labs to implement, particularly in preserved tissue, in part due to the complexity and cost of existing technologies at the level of probe design, tissue pre-treatment, method workflows, limitation to spatially separated targets, image registration, and computational alignment. Accordingly, we set out to create a quantitative FISH method that satisfies at minimum the following criteria: (1) highly multiplexed simultaneous amplification, (2) user control over features like levels of amplification and detection efficiency, (3) a simple and robust workflow, and (4) the ability to detect mRNA and genomic DNA targets.

Recently, we introduced a molecular strategy to program the autonomous synthesis of ssDNA *in vitro* termed ‘primer exchange reaction’ (PER),^40^ which enables the growth of long ss-DNA concatemers from a short (9 nt) DNA primer sequence.^40^ By designing a single PER hairpin sequence to act as a catalytic template, identical sequence domains can be repeatedly appended to nascent single-stranded primer sequences with a strand displacing polymerase. Kinetics of the reaction are controllable via a number of factors, enabling simple but effective control over the lengths of concatemers synthesized *in vitro*. We hypothesized these concatemers could serve as an effective platform for fluorescent signal amplification, as their polymeric structure provides an ideal hybridization target for localizing many fluorescent strands to a single strand scaffold and is reminiscent of the sequences found in branched signal amplification approaches.^15–18^ Furthermore, because PER enables a simple and effective platform to design and synthesize a large number of orthogonal sequences, we reasoned that we could readily implement serial multiplexed imaging strategies based on programmable DNA hybridization and removal^41–43^ into the detection workflow to achieve highly multiplexed imaging.

Here, we introduce a new molecular toolkit termed signal amplification by exchange reaction (SABER) that harnesses the programmability of PER to enhance the functionality of oligo-based FISH probes such as single-molecule RNA FISH probe pools^10^ and highly complex ‘Oligopaint’ probe sets.^44^ Briefly, DNA and RNA FISH probes are first chemically synthesized with primer sequences on their 3’ ends and then concatemerized using PER *in vitro*. Extended probe sequences are then hybridized *in situ* and act as scaffolds to which multiple fluorescent strands can bind. We establish that SABER provides a fast, simple, and inexpensive platform to amplify the signal of both RNA and DNA FISH probes in fixed cells and tissues. We experimentally demonstrate that in different scenarios, SABER can amplify a signal up to 450 fold, can be deployed against up to 17 different targets simultaneously, and can provide high sampling efficiency of target transcripts for puncta detection and cell type identification. We further highlight an application of SABER in tissues by applying a novel 10-plex FISH assay for interrogating the activity and specificity of candidate enhancer elements in mouse retinal tissue via the detection of reporter RNAs, followed by co-detection of reporters and plasmids through a combined RNA/DNA FISH experiment.

## RESULTS

### Design of orthogonally detectable sequences for use in building probe-linked concatemer structures

We recently developed the primer exchange reaction (PER) method for autonomously synthesizing arbitrary single-stranded DNA sequences from short DNA primers.^40^ The method can be used to programmably append specific DNA sequence domains onto growing single-stranded concatemer sequences. One embodiment of the reaction uses a catalytic hairpin and strand displacing polymerase to repeatedly add the same sequence domain onto single-stranded primers, as shown in the PER cycle depicted in Fig. 1A.

**Figure 1:**
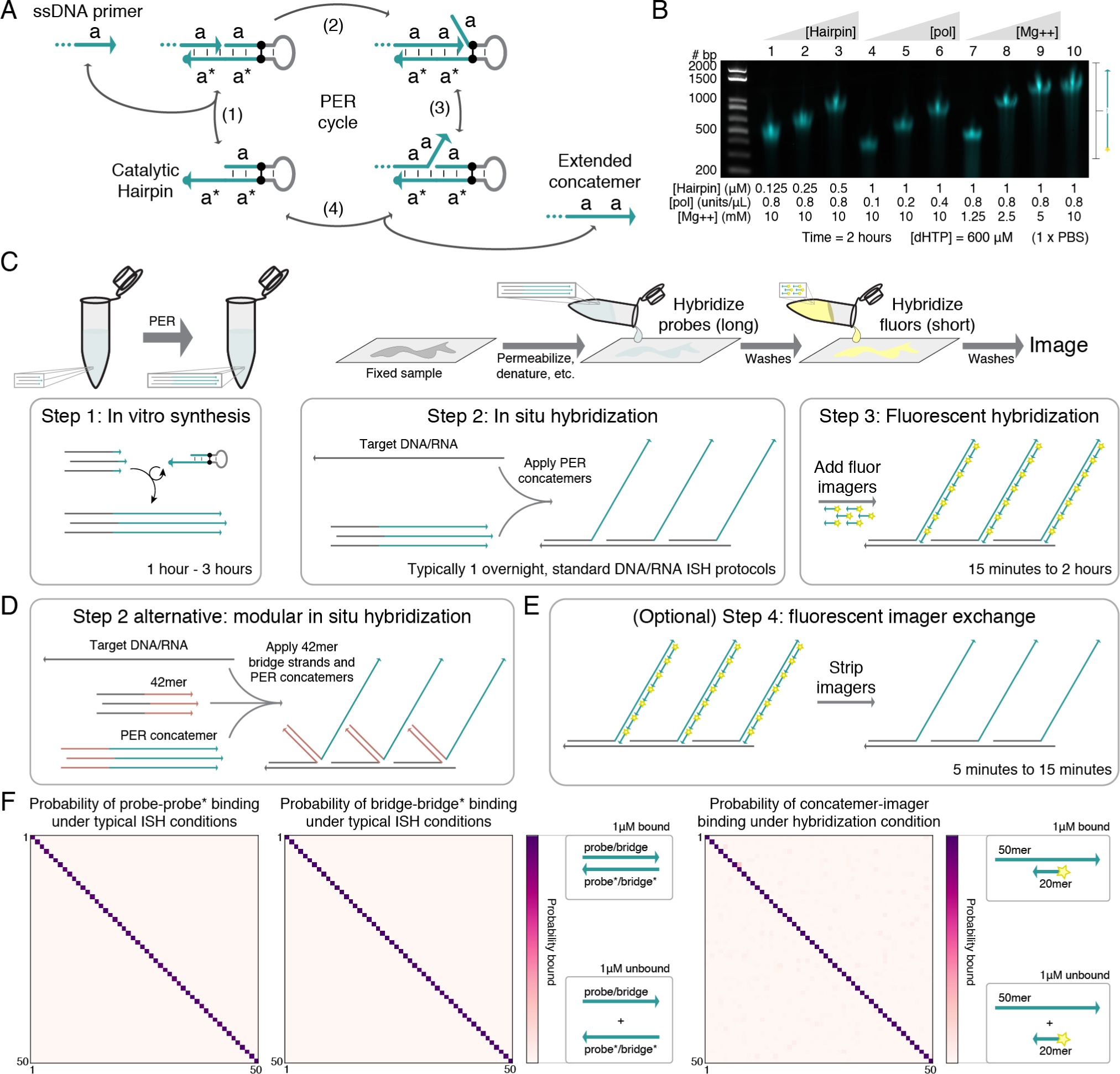
SABER-FISH design and workflow. (**A**) Primer exchange reaction (PER) cycle.^40^ Single-stranded DNA primers are extended by catalytic hairpins. A primer with domain **a** on its 3’ end binds (reversibly) to a catalytic hairpin (step 1) and then gets extended with a new **a** domain by a strand displacing polymerase (step 2). Competitive branch migration (step 3)^45^ displaces the newly extended primer, which can then dissociate (step 4). Primers can undergo the cycle repeatedly, resulting in the generation of long repetitive concatemeric sequences. (**B**) Length programmability of PER concatemers. Hairpin concentration, polymerase concentration, magnesium concentration, and incubation time are all parameters that can be adjusted to control PER kinetics and therefore concatemer length. Time and dHTP (dATP, dTTP, and dCTP) concentration were fixed at 2 hours and 600 µM, respectively. See also Fig. S1A. (**C**) SABER workflow. First, large batches of probe sequences with primers on their 3’ ends can be concatemerized using PERs (see schematic in Fig. S1C). These extensions can then be hybridized to DNA and RNA targets in fixed samples following standard *in situ* hybridization (ISH) protocols. This is followed by a short step that hybridizes fluor-conjugated oligos to complementary concatemers. (**D**) An alternative method uses 42mer ‘bridge’ sequences to couple probes to concatemer sequences. This method allows the same 42mer barcodes with PER concatemers to be re-used for different targets. (**E**) Optionally, fluorescent signal can be rapidly stripped from concatemers under conditions that preserve the underlying probe sequences to stay bound, allowing exchange of imager strands^41, 41^ targeting different concatemers using the same fluorescence channel. (**F**) Orthogonalities of probes, 42mer bridge, and PER sequences. NUPACK^46–48^ is used to evaluate the binding probabilities of complementary and orthogonal sequences. The probabilities of an example set of 50 probes binding to their 50 complementary sequences (left) and of 50 42mer bridge sequences binding to their 50 complementary sequences (middle) under typical FISH conditions (2×SSCT + 50% formamide at 42°C) shows high likelihood of cognate pairs binding (diagonal of matrix) with little cross-talk between non-cognate sequences. A similar analysis for 50 computationally designed PER primer sequences, analyzing binding probabilities of 50mer concatemers to their complementary 20mers under typical hybridization conditions (1×PBS at 37°C) also shows high levels of orthogonality. See Methods section for additional information on sequence design.

We found that the length of repetitive PER concatemers produced in an *in vitro* reaction could be tuned using several different parameters, including polymerase concentration, hairpin concentration, and magnesium concentration (Fig. 1B). Varying the incubation time and nucleotide concentrations can also modulate the length of concatemers (Fig. S1A). We designed the concatemers to contain no G bases, both so that a G-C pair could be used as a PER polymerase stopper in the absence of dGTP and to minimize secondary structure that could inhibit the growth of long sequences. With this simple and effective ability to program the reaction kinetics and therefore length of concatemers *in vitro*, we asked whether this process could be applied in FISH applications as a method for signal amplification by using the concatemers as scaffolds that localize many fluorophores to a single locus.

In signal amplification by exchange reaction (SABER), PER concatemers are extended *in vitro* onto chemically synthesized probes bearing a 9-mer primer (Fig. 1C and Fig. S1B-C). Probes can be generated in bulk quantities, with a user-defined amount applied in hybridization, which is performed using standard DNA or RNA *in situ* hybridization (ISH) protocols. Primary probe hybridization is followed by a short hybridization of fluorophore-labeled ‘imager’ oligos, which bind to the concatemeric sequences. In comparison to methods that generate concatemers *in situ*, the ability to pre-extend large quantities of probe and perform quality control prior to hybridization reduces the time, cost, and variability of the workflow.

A more modular version of this FISH workflow, which uses 42 base pair (42mer) ‘bridge’ sequence domains to hybridize concatemers to FISH probes, is depicted in Fig. 1D (see also schematic in Fig. S1D). For each new target, relatively short probe sequences with 30-50bp homology to their targets can be designed using standard approaches, and then one of 84 designed 42mer bridge sequences can be appended to the end of the sequence. Then, the complementary 42mer sequence can be concatemerized *in vitro* from a PER primer on its 3’ end and co-hybridized together with oligopaint FISH probes. This strategy allows the same bulk sets of extensions to be re-deployed for different targets and samples as needed and further reduces the cost by allowing the same sets of concatemers to be re-used for each different application. Complementary fluorescent ‘fluor’ imagers that have 20 bases of homology to the concatemer are typically used for imaging. Optionally, these 20 nt imager oligos can also be stripped from bound concatemers to reset the signal,^41, 41^ enabling subsequent use of that fluorescence color on a distinct target (Fig. 1E).

Sequence orthogonality in all aspects of the design was considered to ensure robust and specific targeting of fluorescent signal. We use the OligoMiner pipeline^49^ to design orthogonal ‘oligopaint’ probe sequences with homology to targets of interest. By using a series of computational tools, probe sequences are vetted for orthogonality against the relevant target genome, and single-strandedness and melting temperature constraints are used to further filter sequences. Finally, FISH probes are hybridized under conditions close to their melting temperature to increase specificity of binding (Fig. S1E). We can model the probabilities of an example set of 50 such probe sequences binding to each other’s complements using NUPACK^46–48^ and plot the pairwise results in an orthogonality matrix (see Fig. 1F, leftmost plot). Under typical FISH conditions (2×SSCT with 50% formamide at 42°C), probes and their complementary sequences have high probabilities of binding to each other (diagonal), whereas non-cognate pairs of probes and complements have very low predicted binding probabilities in these conditions. A similar design process, but with sequences drawn from blocks of orthogonal sequences,^50^ was used to generate 84 orthogonal 42mer bridge sequences. An analogous orthogonality matrix for 50 orthogonal bridge sequences is depicted in Fig. 1F, middle plot. The full sequence list of 42mer bridge sequences designed can be found in Supplemental Section 8.

One of the major proposed advantages of concatemer-based amplification has been the ability to generate orthogonal sequences that enable simultaneous multiplexed visualization of targets *in situ*. Instead of amplifying each single target iteratively, multiple distinct target signals can be amplified in parallel and visualized simultaneously using spectrally distinct fluorophores. To achieve highly multiplexed imaging using SABER, we needed to design many orthogonal PER concatemer sequences. We previously showed how dozens of orthogonal PER sequence domains can be extended together in a single test tube to reliably create staple strands for a DNA origami nanostructure and how multiple orthogonal PER primers can be concatenated together to form arbitrary sequences.^40^ For repetitive PER concatemers to be orthogonal, it is important that they have high likelihood of binding complementary imager strands while still being unlikely to bind to non-cognate imagers. As with the probe and 42mer bridge sequence modeling, we used NU-PACK^46–48^ to model the probabilities of these on-and off-target interactions for 50 computationally designed PER primers under the desired fluorescent hybridization buffer conditions (1×PBS at 37°C). The pairwise binding probabilities for 20mer imaging sequences to 50mer concatemers are depicted in the rightmost orthogonality matrix in Fig. 1F, pairwise binding probabilities of the sequence sets to each other can be seen in Fig. S1F, and the full set of 50 primer and associated hairpin sequences can be found in Supplemental Section 8.

### SABER effectively amplifies fluorescent signals

We first applied SABER to DNA and RNA targets with known distribution in cell culture samples treated with standard PFA fixatives. First, a DNA oligo with homology to the human telomere sequence was extended to 5 different lengths (conditions E1-E5) using different concentrations of hairpin (Fig. S2B). The fluorescence resulting from hybridization with probes having each of these lengths, and a probe having an unextended sequence with a single binding site for fluorescent imagers (condition U), was visualized using fluorescence microscopy (see Fig. 2A and Fig. S2A). A custom CellProfiler^51^ pipeline was used to identify and quantify puncta within cell nuclei (see Methods section for additional details). Distributions of peak puncta fluorescence values for all conditions were visualized (Fig. 2B, top), and fluorescence fold enhancement was estimated by subtracting background and dividing by the mean of the unextended condition (Fig. 2B, bottom). We estimated 6.2×, 5.0×, 8.6×, 6.8×, and 13.3× fold enhancement for conditions E1 through E5 relative to the unextended (i.e. unamplified) probe. See Methods and Supplemental Section 8 for additional details.

**Figure 2:**
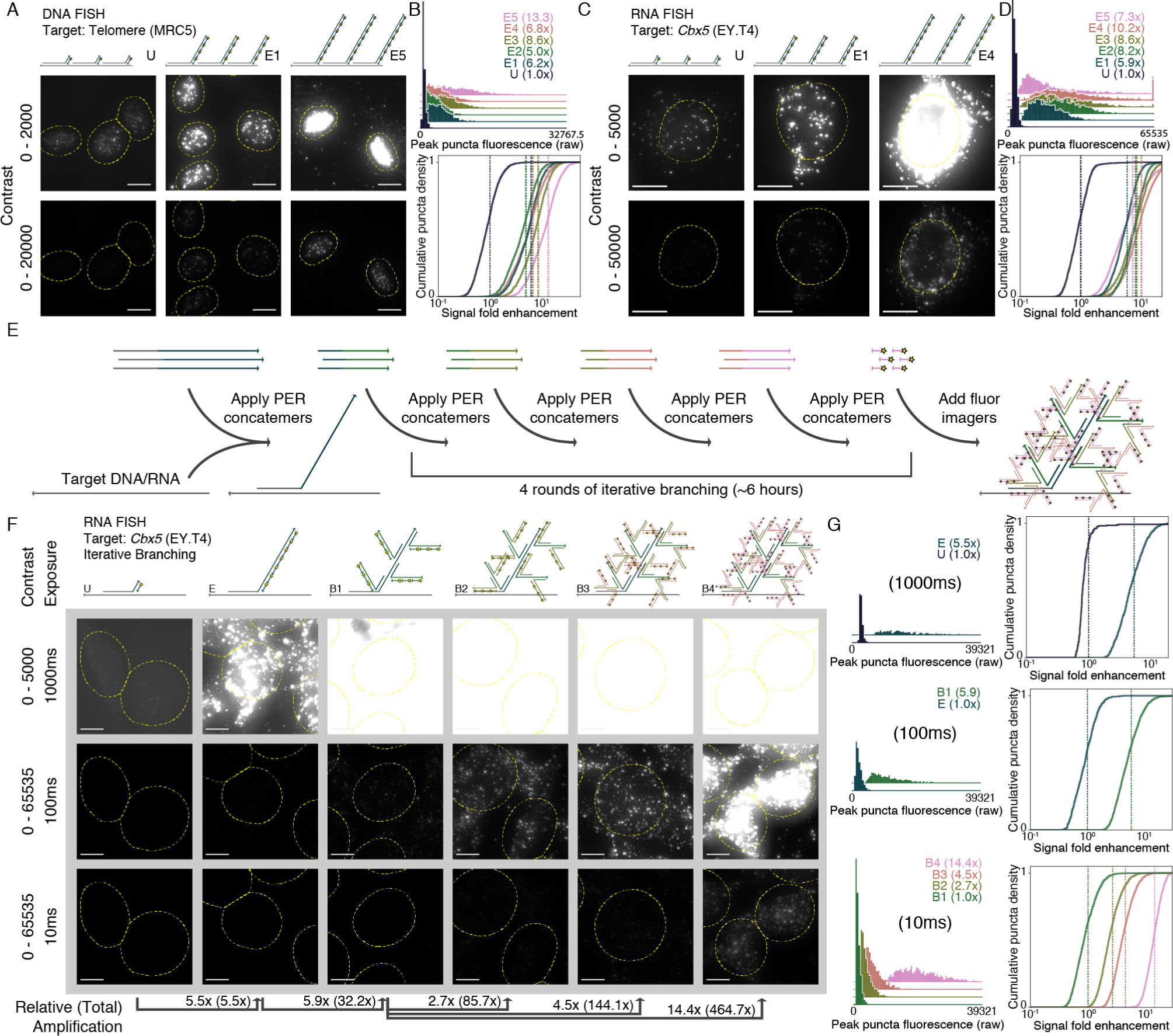
SABER effectively amplifies fluorescent signals. (**A**) Microscopy images for unextended (U) and two increasing lengths of concatemerized probes (E1 and E5) targeting the human telomere sequence are depicted under two contrast levels. See Fig. S2A for additional images and Fig. S2B for a gel showing concatemer lengths. (**B**) An automated CellProfiler^51^ pipeline was used to determine peak fluorescence intensity values for detected puncta in each condition (top). Normalized background-subtracted cumulative distribution functions were plotted to show fold enhancement over the unextended condition (U), with vertical lines depicting normalized distribution means (fold enhancement). See Methods section for additional details. n=1,846-2,190 puncta. (**C**) A similar method was used to visualize unextended (U) and several extension lengths (E1 to E5) of a 122 probe pool targeting the mouse *Cbx5* mRNA transcript. See Fig. S2C for additional images and Fig. S2D for a gel showing concatemer lengths. (**D**) Another automated pipeline that identified puncta within cell bodies was used to quantify amplification in a similar manner to part (B). Distributions show moderate increases in fluorescence for longer extensions, until the longest, which shows a dropoff in signal level. n=1,588-3,279 puncta. (**E**) Multiple rounds of PER concatemer binding were used to dramatically amplify signal further. (**F**) Representative images for samples with unextended probes (U), extended (E), and one through four rounds of additional PER concatemer binding (branching, B1 through B4) are shown under three different exposure and contrast conditions. (**G**) A CellProfiler^51^ pipeline automatically detected puncta, and relative fluorescence was compared as previously but with no background subtraction to determine amplification fold enhancement for extended vs. unextended. In total, amplification fold enhancement over unextended probes (with no background subtraction) was estimated to be 465×. n=262 puncta (unextended), n=717-2,059 puncta (extended, branched). Scale bars: 10 µM.

Next, the process was repeated for a set of 122 probes de-signed to target the mouse *Cbx5* mRNA transcript with 3’ extensions containing a single binding site for a 20mer fluor oligo. Here we employed a relatively large probe set to ensure unam-plified signal could be properly visualized and quantified. The probes were pooled together, extended to five different lengths (Fig. S2D), and visualized (Fig. 2C and Fig. S2C). Puncta within cell bodies were segmented (see Methods), and signals from ex-tended versus unextended probes were compared and quantified (Fig. 2D). The first four extension lengths showed increasing levels of amplification (5.9×, 8.2×, 8.6×, and 10.2× fold en-hancement for conditions E1 through E4), but the longest exten-sion (condition E5) showed a dropoff (7.3×), indicating the im-portance of extension length programmability available through modulation of the parameters described above. In general, our results indicate that extension lengths between about ∼250 and 750 nt provide robust, though not substantially different, levels of amplification.

Multiple rounds of PER concatemer hybridzation can further increase fluorescent signal level by creating branched concate-meric structures^15–18^ (Fig. S2E). First, ISH hybridizes probes with concatemer extensions to their nucleic acid targets. Then, a second round of hybridization binds PER concatemers with 30mers having homology to the concatemer sequences of the primary probe. Following a similar imaging and feature identification pipeline as above, one round of branching amplification was visualized and quantified for several branch sequence lengths targeting *Cbx5* mRNA transcripts (see Fig. S2D). Again, we found amplification levels of 35.5× fold enhancement com-pared to unamplified signal (Fig. S1G). Branching therefore serves as a simple and effective method for further amplifying fluorescent signal.

Multiple rounds of branching can result in even higher lev-els of amplification. Because each hybridized concatemer can serve as a target for many concatemers in subsequent rounds, the level of amplification theoretically scales exponentially instead of linearly with the number of rounds. We implemented four rounds of branching on top of probe concatemers targeting the *Cbx5* mRNA transcript (Fig. 2E). The sample with four rounds of branching (B4) was compared to wells that only underwent three (B3), two (B2), and one round (B1) of branching in addition to extended but unbranched (E) and unextended (U) conditions (Fig. 2F). Each field of view image was taken using three exposure times (10ms, 100ms, and 1000ms) (Fig. 2F). After feature segmentation, max pixel values within identified puncta were quantified only under exposure times where puncta were reliably identified. Probability density distributions for the peak fluorescence values of features under these conditions are shown in Fig. 2G, along with the cumulative density distributions, based on fold enhancement without background subtraction over the lowest amplification condition relevant for that exposure time. In total, signal fold enhancement was estimated to be 32.2×, 85.7×, 144.1×, and 464.7× for one, two, three, and four levels of branching, respectively.

### SABER enables robust transcript detection in tissue

We next asked whether SABER could be used to amplify RNA FISH signal in tissue sections. The mouse retina has been extensively characterized by single cell transcriptomics (scRNA-seq),^53, 53^ providing a useful point of comparison to assess the target specificity and quantifiability of FISH using SABER probes. We first compared unextended probes to PER-extended probes by targeting Rhodopsin (*Rho*), expressed exclusively in rod photoreceptors (Fig. 3A). Here, an exceptionally abundant mRNA was selected to permit visualization of unamplified sig-nal in tissue sections. Fluorescent signal was localized to the photoreceptor layer, and was observed in extended and unex-tended conditions, with 5.2×, 6.4×, 7.0×, and 7.9× fold enhance-ment for increasing extension lengths (conditions E1 through E4) versus unextended probes (condition U, Fig. 3B-C). FA-fixed cryosections cut to 35-40 µm thickness were used for these and subsequent experiments.

**Figure 3:**
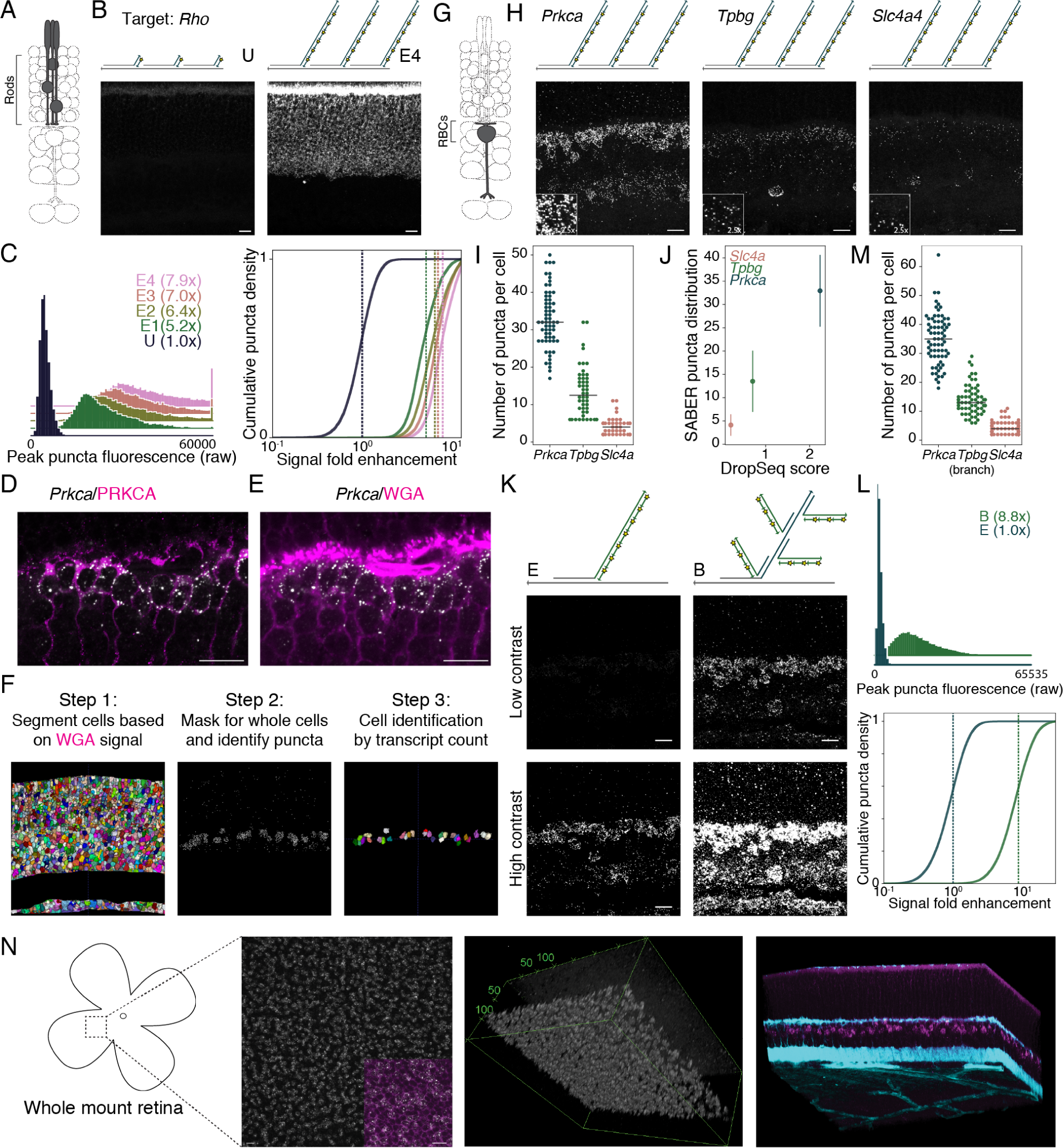
Transcript detection and quantification in retina tissue. (**A**) Schematic representation of retinal cell type (rods) targeted. (**B**) SABER-FISH detection of *Rho* transcript following hybridization with probes conjugated to a single 20-mer fluorescent oligo binding sequence (unextended, U) or with PER-extended probes bearing a concatemer of length ∼650 nt (extended, E4). n=11,159-19,848 puncta. (**C**) Quantification of SABER-FISH signal intensity for *Rho* transcript detection with unextended probe (U) and probes of varying concatemer lengths (E1-E4). See Methods section for additional details. (**D**) Combined *Prkca* transcript detection and PRKCA protein detection by IHC, demonstrating specificity of transcript detection and localization of transcripts relative to the cell boundary. (**E**) Fluorophore-conjugated WGA outlines cell bodies in the retina. (**F**) Analytical pipeline for assignment of puncta to discrete cells in 3D tissue sections using *ACME*^52^ for cell segmentation and custom MATLAB pipeline (*PD3D*) for puncta detection and assignment. See Methods for details. (**G**) Schematic representation of retinal cell type (rod bipolar cells, RBCs) targeted. (**H**) SABER-FISH detection of transcripts for three genes with differential expression levels and highly enriched expression in RBCs relative to other bipolar cell types. (**I**) A swarm plot of SABER puncta per cell is shown with a median line overlay. n=45-63 cells. (**J**) Quantification of RBC marker transcripts recapitulates relative abundance of transcripts (average number of transcripts per cell) observed from >10,000 single RBCs profiled in a Drop-seq dataset.^53^ Standard deviations of SABER puncta per cell are shown. (**K**) Addition of branches provides a large increase in signal intensity while maintaining specific labeling patterns. (**L**) Quantification of the increase in signal intensity provided by the addition of branches to *Prkca* detection. n=29,818-35,330 puncta. (**M**) Quantification of transcripts detected using branched probes. n=65-78 cells. (**N**) Whole mount retina detection of *Grik1* transcript. Schematic shows flat mount retina preparation used for staining and imaging. (Left) Maximum intensity projection of *en face* view in the inner nuclear layer, inset shows WGA counterstain (magenta); (Middle) ImageJ^54^ 3D Viewer rendering of a Z-stack that matches the Z-dimensions of the retina, axis labels in µms; (Right) ImageJ^54^ Volume Viewer rendering showing *Grik1* expression (magenta) and WGA signal visible in the plexiform layers and blood vessels (cyan). Scale bars: 10 µm.

We next tested the performance of SABER probes in the detection of lower abundance transcripts, choosing rod bipolar cells (RBCs), a single type of bipolar interneuron that has been extensively profiled by scRNA-seq,^53, 53^ as a test popu-lation. Specificity of FISH was confirmed by co-detection of the *Prkca* transcript and PRKCA protein, an established RBC marker (Fig. 3D and Fig. S3A). The ability to quantify detected transcripts per cell is important to assess the performance of SABER, and for subsequent applications; therefore, a generalizable and unbiased method for defining cell boundaries is highly relevant. We found that fluorophore-conjugated wheat germ agglutinin (WGA) provides an effective label to outline all cells of the retina and is compatible with SABER-FISH (Fig. 3E), enabling 3D cell segmentation using *ACME*,^52^ an open-source software for membrane-based watershed segmentation (Fig. 3F). A Laplacian of Gaussian method was used to localize fluorescent SABER puncta in 3D, which could be assigned to cells based on their location relative to segmented cell boundaries. Segmented cells can then be assigned cellular identities based on both marker gene transcript counts and laminar position in the tissue (Fig. S3B-D and see Methods section for further details). While WGA-based segmentation is limited by the inability to resolve neuronal processes, this limitation also applies to dissociated single retinal cells, and it is therefore a relevant method for use in comparisons to scRNA-seq.

From the described set of RBC-expressed genes,^53^ we selected three transcripts for quantification that are highly enriched among RBCs (Fig. 3G) and that are expressed at low (*Slc4a*), moderate (*Tpbg*), or high (*Prkca*) levels (Fig. 3H-I and Fig. S3A). We found that relative transcript abundance for these genes in RBCs as detected by SABER-FISH closely paralleled the relative abundance observed by Drop-seq (Fig. 3J). Sampling of transcripts by SABER was approximately 15-fold higher than observed for Drop-seq-profiled cells, where cells had been sequenced to an average depth of 8,200 reads per cell for comprehensive classification of bipolar cell types (50-fold deeper sequencing of Drop-seq libraries improves transcript detection probability by up to ∼2-fold^53^). Transcript quantification for these probes remained stable when branch detection was employed (Fig. 3K), and we observed an 8.8× fold increase in signal intensity using a single round of branching compared to simple extension (Fig. 3L-M). The application of branching permitted robust detection of transcripts with a probe set composed of just 12 probes (Fig. S3F). A concern in the application of pre-extended probes to tissue is the ability of long DNA strands to penetrate. We tested SABER-FISH in FA-fixed flat-mounted retinas (∼150 µm thickness), modifying the tissue section protocol to have longer incubation and wash times, and observed effective labeling of mRNA in bipolar cells of the inner nuclear layer (Fig. 3N).

### SABER enables spectrally multiplexed imaging

Probes that target distinct transcripts can be visualized simultaneously by appending orthogonal concatemer sequences detectable by imager oligos with distinct fluorophores. Three repetitive regions of the mouse chromosome - major satellite, minor satellite, and telomere - were visualized simultaneously in mouse retinal tissue by using this approach (Fig. 4A), permitting observation of the distinctive chromatin architecture of rod photoreceptors.^56^ Another three-color visualization was performed in human metaphase spreads and interphase cells to target three co-localized positions on Chromosome 1 (Fig. 4B). In total, 18,000 probes targeting 3.9 Mb region were mapped to three colors, which all co-localized as expected. Intronic and exonic sequences were also separately detected for the *Dll1* mRNA transcripts in developing retina (Fig. S4A), a distinction that is useful as a method to probe transcription kinetics.

**Figure 4:**
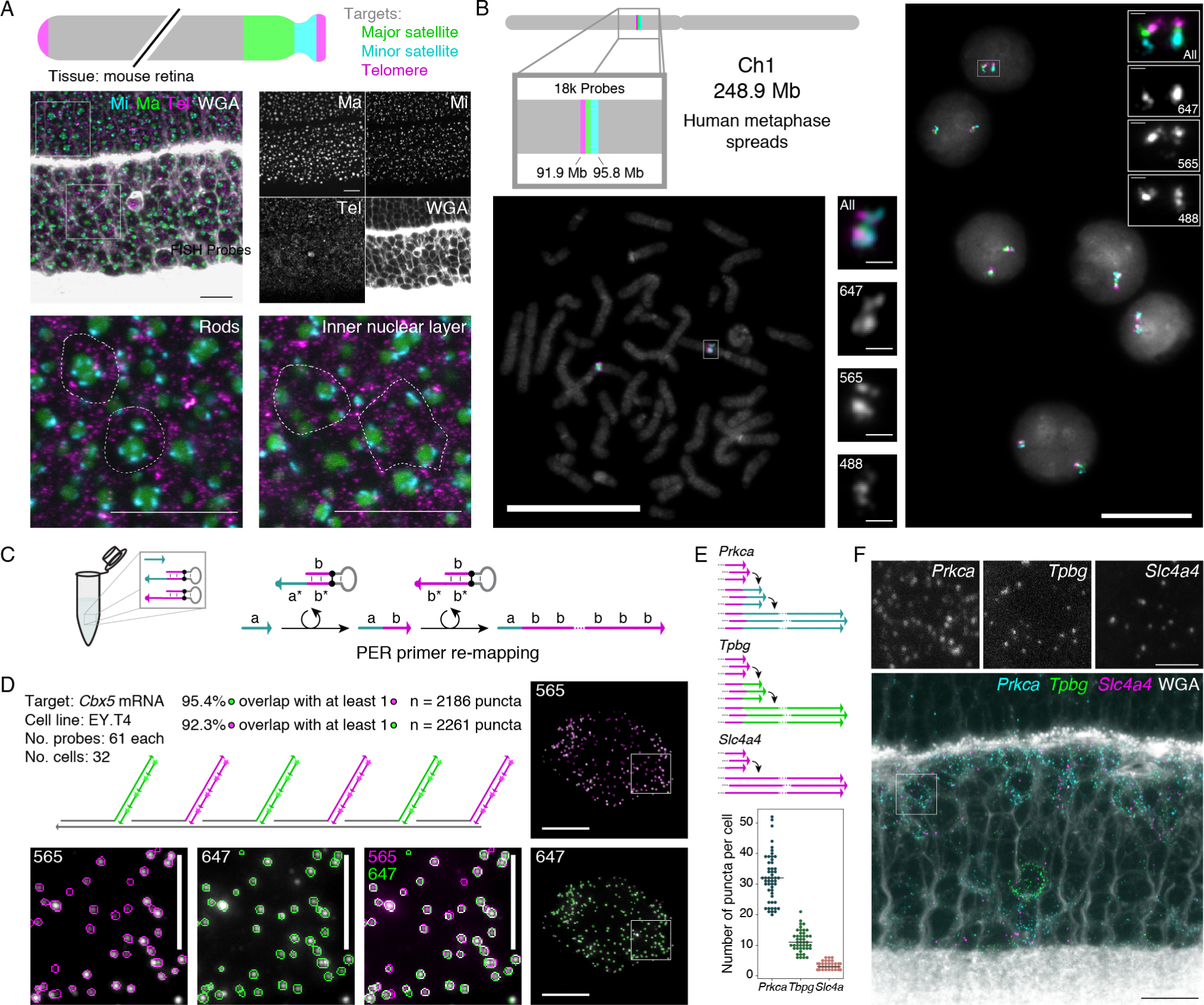
SABER enables spectrally multiplexed imaging. (**A**) Multiplexed SABER-FISH in mouse retina. Mouse major satellite, minor satellite, and telomere chromosomal regions were detected with orthogonal SABER concatemers and visualized simultaneously. Magnified views show distinct organization of these chro-mosome regions in rods compared to inner nuclear layer cells. Dashed outlines are manually drawn approximations of the cell boundary based on WGA staining. Scale bars: 10 µm. (**B**) Multiplexed SABER-FISH on metaphase spreads. Three adjacent positions on human chromosome 1 were visualized using the 42mer bridge strategy depicted in Fig. 1D, and co-localization of targets was validated in metaphase spreads and interphase cells. Scale bars: 20 µm. (**C**) Primer re-mapping with PER. Primers (e.g. with domain **a**) can be concatemerized with a different repetitive sequence (e.g. primer domain **b**) through the use of a stepwise PER hairpin (e.g. that appends **b** to **a**)^40^ and a standard repetitive hairpin. The primer re-mapping and concatemerization can be done autonomously in a single reaction tube. (**D**) Single molecule co-localization. Primer re-mapping was used to map two halves of the *Cbx5* probe pool (used in Fig. 2) to two new PER primers, and two-color co-localization was then visualized and quantified using another custom CellProfiler^51^ pipeline (see Methods and Fig. S4B). In total, 92.3% of identified puncta in the 565 channel ovelapped with puncta in 647, and 95.4% of identified puncta in 647 overlapped with puncta in the 565 channel. Scale bars: 10 µm (cells), 5 µm (panels). n=2,186-2,261 puncta. (**E**) Primer re-mapping and 3-color visualization in retina tissue. The *Tpbg* and *Prkca* probe sets from Fig. 3 were re-mapped to two new primers to enable simultaneous visualization and quantification of *Prkca*, *Tpbg*, and *Slc4a* transcripts. Representative images are depicted in panel (**F**) and puncta distribution compared to their Drop-seq values^53^ are plotted in Fig. S4C. n=38-52 cells. Scale bars: 10 µm, 2.5 µm (overlay).

We previously showed how PER cascades can be pro-grammed to autonomously undergo differential synthesis path-ways by modulating the hairpins present in solution.^40^ This flex-ibility to program sequences allows us to take existing probe sets and change the sequence of the PER concatemer synthesized onto them. Fig. 4C shows an example of how a primer **a** can be mixed with two hairpins to produce a concatemer with repeats of sequence **b**. The first of two hairpins appends the **b** sequence 3’ from the **a** sequence, and then a second hairpin repeatedly adds the **b** sequence to generate a concatemer with a different PER primer sequence than the original one.

This re-mapping strategy is useful for re-mapping existing probe sets with conflicting (identical) 3’ primer extensions to orthogonal sequences, and is simply achieved by including a primer switching hairpin in the *in vitro* extension reaction. The *Cbx5* probe set (used in Fig. 2) was split into two pools, and each pool was re-mapped to a new primer sequence. This en-abled a two-color visualization of *Cbx5* transcripts (see Fig. 4D), where half of the probes were mapped to the 565 channel and the other half were mapped to 647. A custom CellProfiler^51^ pipeline was used to identify transcript puncta in each channel and compare the number of identified features that overlapped features in the other channel. We found that 92.3% of puncta identified in the 647 channel overlapped with puncta in the 565 channel, and conversely that 95.4% of puncta identified in the 565 chan-nel overlapped with puncta in the 647 channel. These numbers further indicate that SABER-FISH probes can enable detection of a large fraction of available transcripts at the single-molecule level. We also evaluated primer re-mapping for two of the three RBC-expressed genes evaluated in Fig. 3 (*Prkca* and *Tpbg*) to simultaneously detect the three transcript species originally syn-thesized with identical primers (Fig. 4E-F and Fig. S4C).

### SABER enables fast exchange for highly multiplexed se-quential imaging in cells and tissues

Higher levels of multiplexing can be achieved by iteratively de-tecting nucleic acid targets.^41–43^ One approach^41, 41^ uses for-mamide to rapidly destabilize short fluorescent imager strands without destabilizing the primary probe, permitting re-use of spectral channels. By modeling the melting temperatures of 20mer imagers, 42mer bridge sequences, and FISH probe sequences (Fig. S5A), we determined that 50-60% formamide in 1×PBS should effectively and rapidly de-stabilize imagers without significantly affecting the underlying probe and 42mer bridge sequence stability.

We tested this oligo exchange approach in retinal tissue. Neural tissues typically display high cell type heterogeneity, requiring multiplexed detection methods for comprehensive identification of cellular populations. We aimed to detect all seven major cell classes in the retina (Cone, Rod, Horizontal, Bipolar, Amacrine, Ganglion, and M¨uller glia cells) using SABER probes against established markers. A pool of seven primary FISH probes were hybridized simultaneously, and detected in three sequential rounds of secondary fluorescent oligo hybridization (Fig. 5A). Imager oligo exchange occurred effectively in tissues, permitting re-use of spectral channels. We observed the expected laminar separation of the cell classes by visual inspection and by quantification of the spatial distribution of marker-positive cells (Fig. 5B-D). Following serial FISH probe detection, protein epitopes for Prkca and Calretinin were still detectable by IHC and tissue integrity appeared well-preserved, with sublaminae of the inner plexiform layer (IPL) clearly discernible. We also found that DNAse I and Exonuclease I enzymes could be used to strip both primary SABER probes and fluorescent imagers in tissue, while preserv-ing mRNA integrity, as assayed by the ability to perform a sec-ond round of mRNA detection (Fig. S5B). Therefore, high mul-tiplexing using SABER is achievable by both a large selection of concatemer sequences and the ability to recycle concatemer sequences through primary probe digestion and rehybridization.

**Figure 5:**
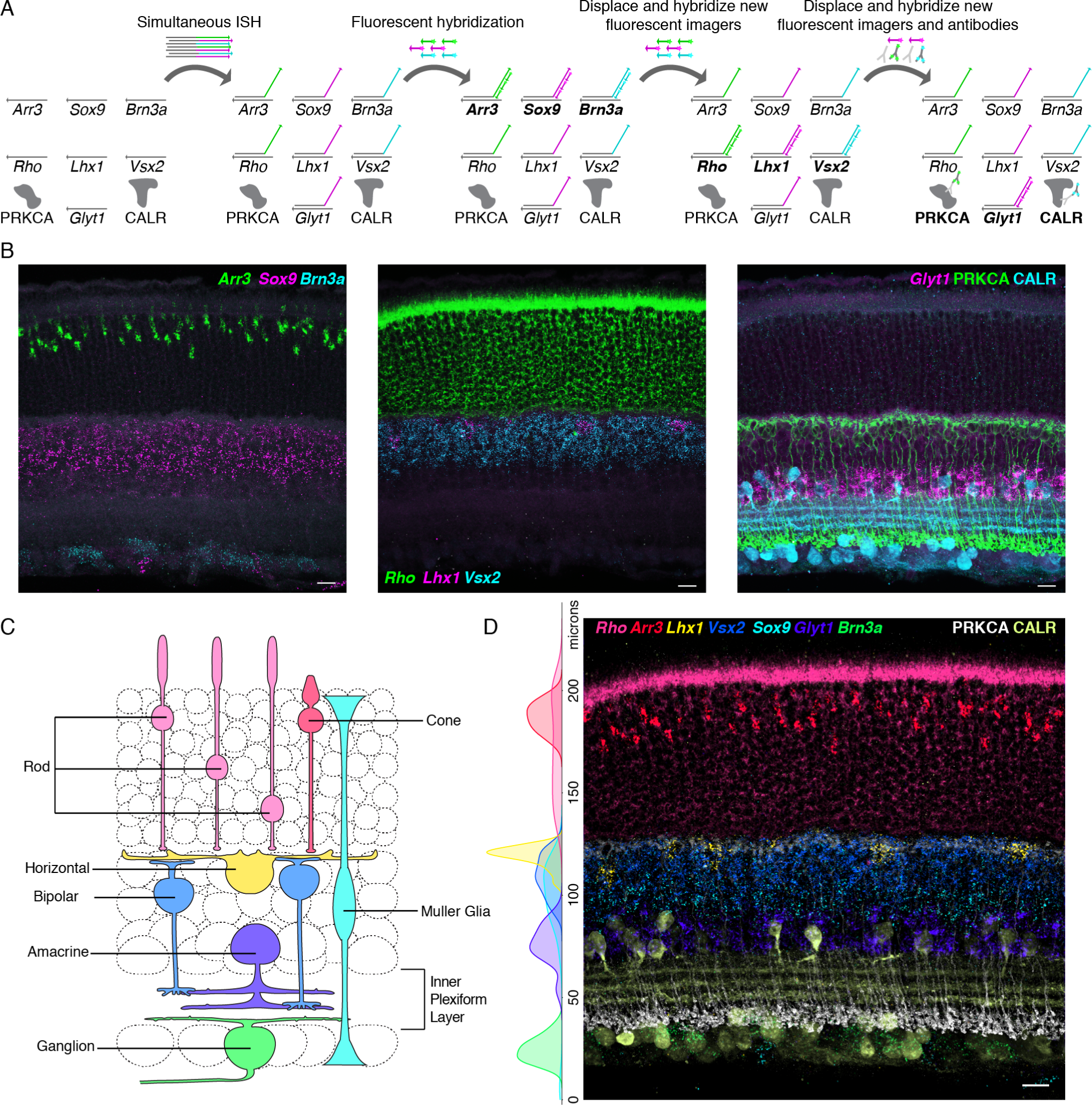
Serial SABER-FISH for detection of cell types in retina tissue. (**A**) Experimental design for serial FISH detection of 7 transcripts and IHC detection of two protein targets. (**B**) Three FISH detection cycles for identification of 7 retinal cell populations. IHC was performed after the 3rd FISH detection. (**C**) Schematic representation of the retinal cell classes detected. (**D**) All nine channels overlaid following puncta thresholding, with additional tissue autofluorescence background subtracted by a gaussian filter masking channels around detected puncta. See Methods section for additional details. Side plot: Marker positive segmented cells plotted by distance from the inner limiting membrane display the expected laminar cell type organization. n=649 cells. Scale bars: 10 µm.

We also evaluated exchange imaging with SABER in human metaphase spreads and interphase cells (Fig. 6). One of the advantages of using SABER with signal amplification is the ability to reduce incubation times by taking advantage of the improved reaction kinetics conferred by the presence of many binding sites, since not all binding sites need to be saturated to be able to discern signal. In total, 17 colors (6 hybridizations) were imaged in 7 hours, including stripping, re-hybridization, field of view finding, and z stack imaging times. These 17 colors targeted seventeen 200 kb regions spread along the Human X chromosome (Fig. 6A). Single z slices with signals in focus (metaphase) or maximum projections of the z stacks (interphase) were then automatically aligned based on their DAPI signals (Fig. 6B-C and Fig. S6A). Overlays of all colors onto DAPI signal can be seen in Fig. 6C. Metaphase spreads validate the directional coloring of targets, and interphase cells depict X chromosome territories within their nuclei.

**Figure 6:**
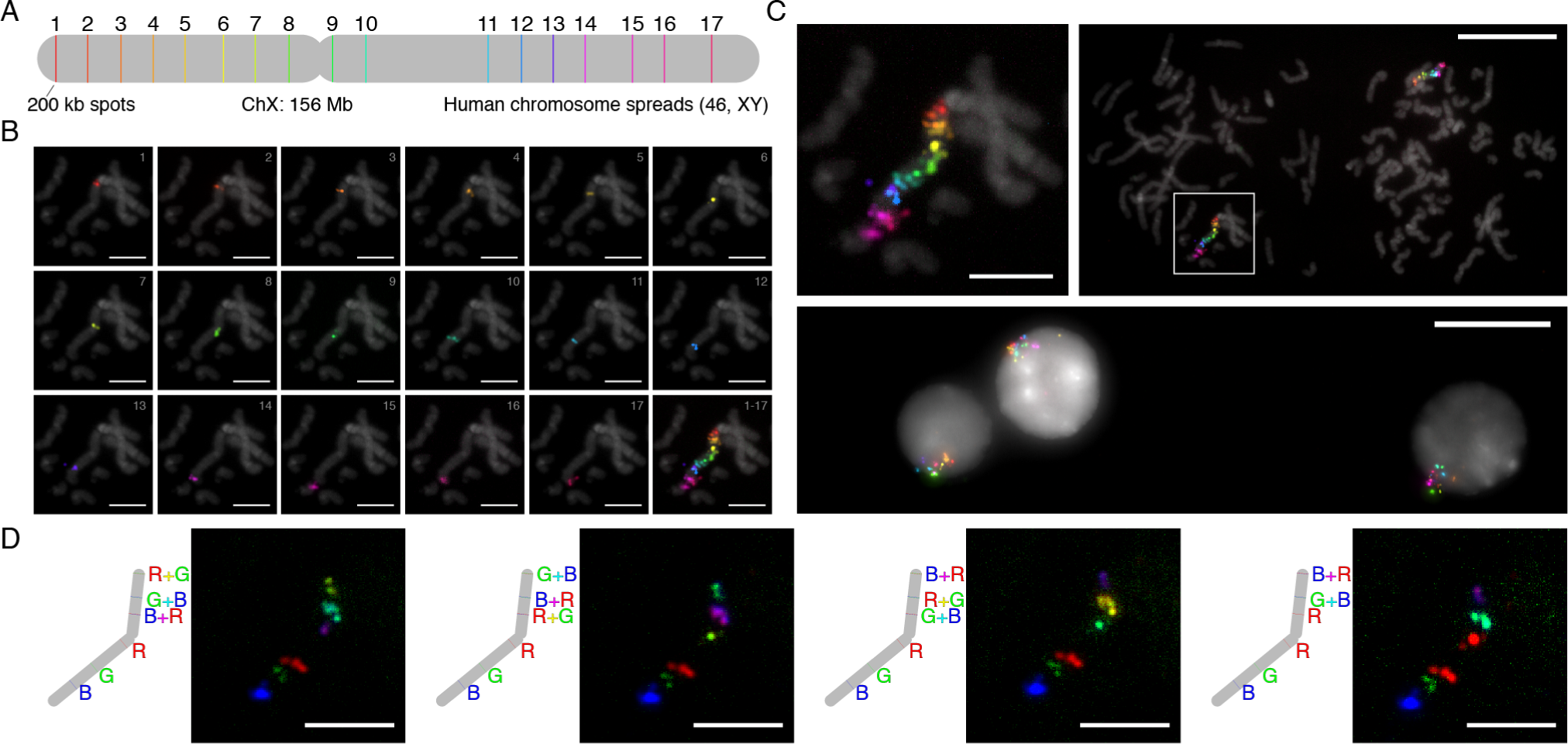
Rapid, highly multiplexed sequential imaging using serial SABER-FISH. (**A**) Schematic of targeted X chromosome. 17 regions along the human X chromosome (width to scale) were targeted with probe pools generated using the OligoMiner pipeline.^49^ Each set of probes per spot had different 42mer barcode sequences appended to their 3’ ends (see Fig. 1d). This allowed seventeen 42mer bridge sequences concatemerized with 17 different PER primers to be co-hybridized during an overnight *in situ* hybridization step. (**B**) Individual color channels on DAPI. A total of 6 hybridizations, targeting 3, 3, 3, 3, 3, and 2 spots respectively, that took course over a single day were used to image the 17 colors. Channels were contrasted and pseudocolored to their respective rainbow hues. (**C**) 17 color overlays on DAPI. The representative metaphase from part (B) is shown overlaid on DAPI at two length scales (top left, top right image). Interphase cells showing the X chromosome territories were also captured (bottom image). (**D**) Combinatorial and 6-color SABER imaging. As a step toward increasing multiplexing with SABER amplification further, we demonstrated mapping six of the spots on the chromosome to 4 different 6-color combinations. Scale bars: 5 µm (spreads), 20 µm (fields of view).

Several combinatorial approaches^32–35,37,39^ have recently been presented as frameworks to dramatically increase the number of colors (targets) that can be visualized using nucleic acid sequence barcoding of spatially separated targets. We explored the viability of such an approach for future SABER applications. To test this, we took the same metaphase sample used in Fig. 6B-C and applied new fluorescent hybridizations. Each hybridization targeted the same set of 6 regions, mapping them to each of three colors in addition to the pairwise combinations of the three colors, achieved by co-hybridizing complementary imager oligos conjugated with different fluorophores. This process was repeated four times, each with a different 6-color mapping (Fig. 6D). In each case, only the color or colors expected at each position were observed. By expanding the number of colors that can be visualized at once (to 6), and allowing each target to have a different color code permutation, this approach could in theory enable 6^4 = 1,296 colors to be imaged with just four hybridizations (∼4 hours). If the process is repeated for 6 hybridizations (7 hours for the data shown in Fig. 6B-C), up to 6^6 = 46,656 distinct targets could be visualized. This validation of SABER for compatibility with combinatorial imaging strategies is an important step for future, even more highly multiplexed and amplified detection.

### Application of SABER for quantitative *in situ* reporter assay

We next investigated whether the ability of SABER to provide multiplexed, amplified detection of both RNA and DNA sequences in tissues could be applied to reporter assays involving the introduction of exogenous DNA elements. An ideal reporter assay would permit simultaneous quantification of the expression of reporter molecules, the number of introduced DNA constructs encoding the reporters, and the expression of endogenous markers. RNAs are well-suited to function as reporters of cis-regulatory module (CRM) activity as (1) mRNAs provide a more direct read-out of transcriptional activity than proteins and (2) single RNA molecules are discretely quantifiable with standard microscopy. We applied SABER to the detection of reporter RNAs generated from the introduction of non-integrating plasmid reporter constructs, which permit investigation of the activities of isolated CRMs. First, six sequences encoding reporter RNAs detectable by similarly sized probe sets were cloned downstream of a minimal TATA promoter. Each re-porter was validated independently for the ability to report specific patterns of enhancer activity by upstream insertion of a validated Vsx2 enhancer that drives expression in bipolar cells,^57^ followed by electroporation *in vivo* into the retina (Fig. S3A-B).

We applied this reporter set to evaluate the behaviors of previously uncharacterized CRMs using a 10-plex SABER-FISH experiment. Specifically, we searched for regulatory elements that might drive transcription in subpopulations of bipolar cells, permitting these cells to be specifically labeled and, ultimately, manipulated. We selected for investigation candidate CRMs in the vicinity of the gene *Grik1*, a kainite-family glutamate receptor subunit with strong and enriched expression in most OFF bipolar cells (Types 2, 3a, 3b, 4),^53^ as few genetic tools exist to specifically label this population *in vivo*. Candidate CRMs were identified by inspection of retina chromatin accessibility to DNAse I^58^ in a genomic interval proximal to the transcription start site (Fig. 7A). Six candidate DNA sequences (CRMs 1-6) were amplified from the genome and inserted independently upstream of distinct reporters, and the reporter set was introduced as a pool into the postnatal retina (Fig. 7A-B). In order to evaluate the cell types in which CRMs were active, we used SABER to detect all six reporters as well as four markers of cell types accessible by postnatal retina electroporation (*Grm6*: ON bipolar cells; *Glyt1* and *Gad1*: amacrine cell; *Rho*: rods)^59^ (Fig. 7C-D). Three of the six reporters displayed activity, with one reporter, corresponding to CRM-4, selectively expressed in cells that express *Grik1* (Fig. 7D-E and Fig. S7C-D). A second reporter (CRM-1) showed abundant and specific expression in rods, despite a lack of *Grik1* expression in this population (Fig. 7D-E). To confirm that this assay faithfully reports CRM activity we used a canonical protein-based reporter technique, cloning CRM-1 and CRM-4 into a GFP-expressing reporter. As expected, following electroporation, CRM-1-GFP labeled rods, and CRM-4-GFP labeled bipolar neurons with axonal elaboration in the upper half of the inner plexiform layer, a distinguishing morphological property of OFF bipolar cells (Fig. 7F).

**Figure 7:**
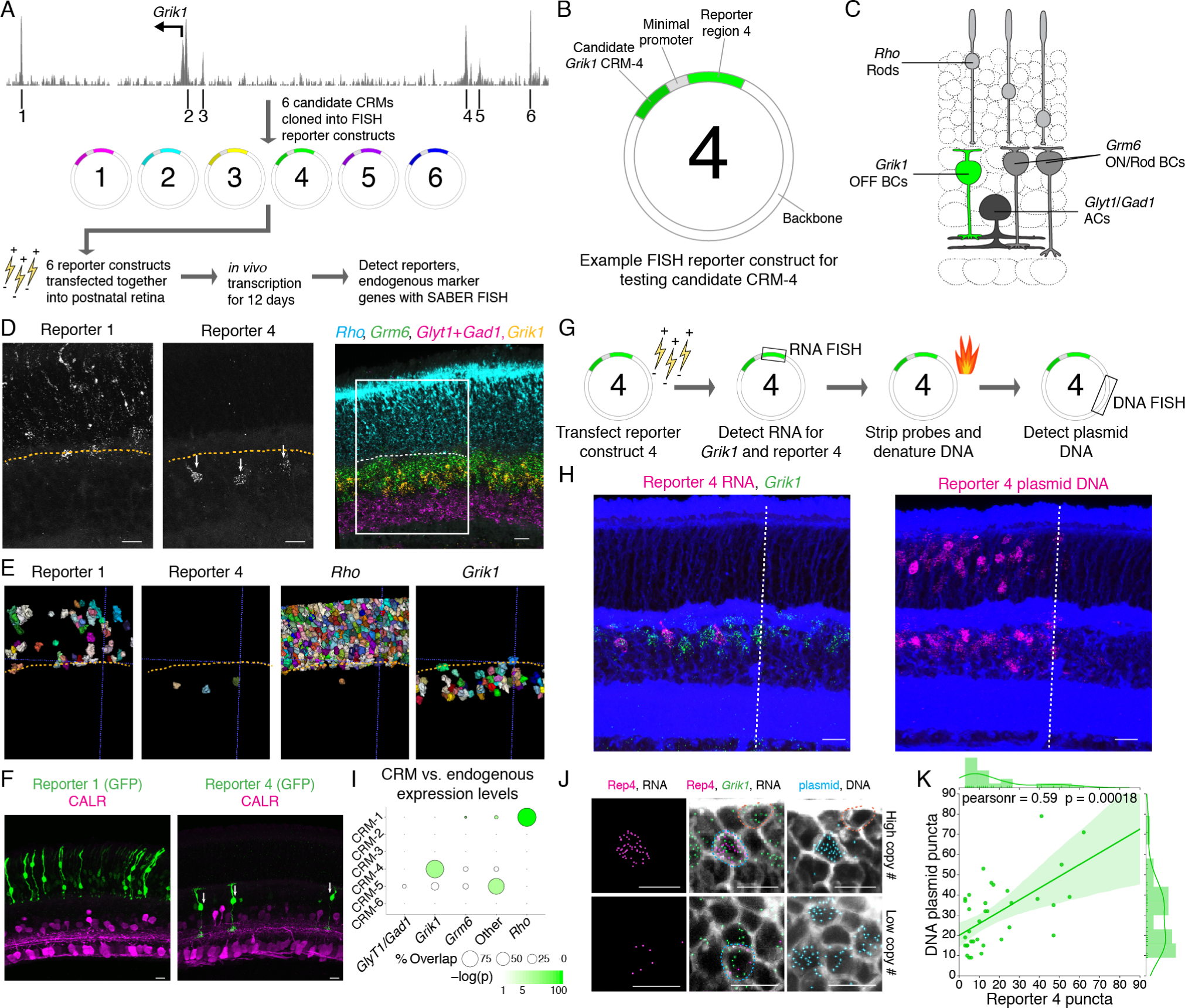
SABER-FISH enables detection of *in vivo* RNA reporters for enhancer activity analysis. (**A**) DNAse I hypersensitive regions in the vicinity of the *Grik1* start site and workflow for the reporter screen experiment. (**B**) Representation of key components of the reporter plasmid. (**C**) Schematic representation of neuronal cell types electroporated in the postnatal retina. *Grik1* expression distinguishes OFF bipolar cells from ON bipolar cells (*Grm6*+). (**D**) Representative images of two expressed reporters (single channel) and four endogenously expressed genes. Dashed line indicates the approximate position of the outer plexiform layer. Box indicates the area of magnification shown for the single channel reporter expression images. (**E**) Relevant reporter and endogenous gene-expressing cells displayed following cell segmentation. (**F**) Expression of GFP driven by CRM-1 and CRM-4 following retina electroporation. Rods (left panel) are identifiable by position in the outer nuclear layer, and OFF bipolars (right panel, arrows) are identifiable by bipolar morphology and lamination in the upper layers of the inner plexiform layer, labeled by Calretinin (CALR). (**G**) Experimental design for reporter RNA and DNA sequential detection. (**H**) Representative image of an electroporated retina with detection of Reporter 4 RNA, *Grik1* RNA, and plasmid DNA. Dashed line indicates the approximate location of the electroporation patch boundary. (**I**) Quantification of the percent of reporter positive cells that are positive for each marker probed. ‘Other’ refers to expression in cells not positive for any marker tested, and may include M¨uller Glia and type 1 bipolar cells. Dot size corresponds to the percent of CRM-positive cells that are positive for each endogenous marker. Dot color reflects the p-value for a hypergeometric test plotted on a logarithmic color scale. (**J**) Magnified images showing detected Reporter 4 RNA puncta in cells with *Grik1* expression and plasmid DNA (blue outline) but not in *Grik1*+ cells lacking plasmid DNA (orange outline). n(cells): n(CRM-1)=440; n(CRM-4)=18; n(CRM-5)=25. (**K**) Quantification of detected Reporter 4 RNA puncta plotted against detected plasmid DNA puncta. n=35 cells. Scale bars: 10 µm.

Finally, we investigated whether SABER could be used to co-detect reporter-encoding plasmids and reporter transcripts. Commonly used methods for dense introduction of exogenous DNAs *in vivo* (e.g. electroporation, AAVs, cationic lipids) can result in a broad distribution of DNA copy number per cell, with cell type-specific biases in transfection rates. This variability can impede assessment of CRM activity (reporter transcripts per reporter DNA). Moreover, in many tissues, cell types are diverse, and the relative abundance of cell types is highly variable. For instance, rods comprise ∼70% of all mouse retinal cells. Measurements of enhancer specificity must account for cell type abundance of transfected cells, and this may not be rigorously achieved by co-transfection of ‘ubiquitously expressed’ (e.g. CMV, Ubc-driven) reporters; these in fact frequently display biased expression between cell types and are not guaranteed to be proportionally co-transfected. To solve this problem, we applied co-detection of the reporter RNA and the plasmid DNA backbone to determine transfected cell type distributions and plasmid load. The CRM-4 reporter was singly electroporated into the retina, followed by detection and quantification of reporter RNA, *Grik1* transcript, and a 2.8kb region of the plasmid backbone (Fig. 7G-H) in the same electroporated cell populations. Using endogenous marker expression and cell position we calculated cell type abundances, determining that 9% of electroporated cells were *Grik1* positive. A hypergeometric test was used to evaluate the probability of observing the empirically determined positive patterns for each CRM, yielding highly significant p-values (Fig. 7I; p value of 1.27×10^−125^ for CRM-1/*Rho* and p value of 1.03×10^−15^ for CRM-4/*Grik1*). We also investigated the relationship between number of plasmids and number of reporter transcripts, and observed a significant correlation (Pearson correlation coefficient of 0.59, p value of 0.00018) that is not observed when comparing plasmid copy number to endogenous *Grik1* expression (Fig. 7J-K and Fig. S7E-F). Variation in transcript number per plasmid may represent selective silencing of plasmids in particular cells, or differences in CRM-4 activity between distinct *Grik1*-expressing OFF bipolar cell types.

## DISCUSSION

The Primer Exchange Reaction (PER) method is a versatile tool for creating user-defined assembly of short sequences using a catalytic hairpin structure. Here we show the application of the telomerase-like mode of PER to achieve enhanced detection of FISH probes. Using a catalytic hairpin bearing identical sequences in the primer binding and copy regions, PER effectively conjugates concatemers of user-defined length to sm-FISH and Oligopaint-style tiling probes. These concatemers provide a scaffold for concentrating fluorescent signal via secondary hybridization, a method we name ‘SABER.’ Through the design of a large array of orthogonally detectable concatemer units we demonstrate the strength of this approach for flexible probe design. (1) Amplification. SABER probes can be greatly amplified for signal detection through the introduction of secondary branches. (2) Modularity. Probes are freely combined for orthogonal detection independent of the intitially syn-thesized primer sequence via bridges or ‘re-mapping’ hairpins that convert the primer sequence. (3) Multiplexing. SABER can achieve high levels of multiplexed, direct, non-combinatorial, detection of RNA and DNA targets, using serial application of fluorescent oligos.

Combined with our recent improvements in the design of on-target probe binding to genomic sequences,^49^ we applied SABER probes in the detection of nucleic acid targets in cells and tissue. SABER proved to be practical to apply in terms of cost and workflow. Probe sets and hairpins could be synthesized without costly base modification or purification methods. Probe sets could be amplified in bulk using PER, allowing enough material for dozens of experiments to prepared in one step and avoiding the need for long amplification steps *in situ*. The combined cost of all oligos and enzymes is currently estimated to be less than $5 per target per experiment (120 µL ISH solution) even in our most expensive conditions, and it is likely this could be significantly reduced with additional optimization and bulk pricing.

Using a single round of branching, we observed strong detection of mRNA transcripts with probes generated from sets of only 12 oligos that were less than 50 nt in length. The protocol requires a single enzymatic step performed *in vitro*, which can be applied to generate probes in bulk, further reducing cost. *In vitro* probe extension has the added advantage of reducing workflow time compared to concatemer generation in tissue, and permits the user to assess the quality (lengths, extension efficen-cies) of concatemers before tissue application. SABER gives effective signal amplification on minimally pre-treated tissues using standard hybridization protocols. We effectively labeled thick tissue sections and whole retinas without the application of tissue clearing or gels. SABER-FISH can be deployed in a straightforward and affordable manner by a common laboratory that already prepares tissue or cells for ISH, and which has access to a conventional confocal microscope.

The analytic pipeline demonstrated here is similarly straight-forward to use. 3D cell segmentation combined with SABER permitted quantification of transcripts on a single cell level. The relative abundance of transcripts closely correlated with measurements from thousands of single cells profiled by Drop-seq. SABER is therefore well suited to accompany scRNA-seq in transcriptomic studies. Sparse sampling of transcripts by scRNA-seq, particularly in the application of droplet-based methods,^60, 60^ can result in the inability to distinguish heteroge-nous expression from low expression. The high efficiency of mRNA detection enabled by SABER can improve quantification of variation in transcript abundance in such cases. The loss of spatial information from tissue dissociation and resulting inability to co-assay other cellular properties such as morphology and activity, is an additional limitation in scRNA-seq approaches. SABER, which is compatible with IHC, can be used to link scRNA-seq-defined populations to positions in tissue such that morphological stains, or labels that permit post-hoc identification of recorded cells, can be integrated with cell type identification. *In situ* transcriptomic methods (STARmap,^39^ seqFISH,^32^ merFISH,^33^ FISSEQ^38^) have been developed to address the loss of spatial information inherent to scRNA-seq. For hybridization-based spatial transcriptomic methods, SABER probe design may provide improvements in signal intensity, sampling efficiency, and ease of application. We also show that combinatorial probe detection is possible with this approach. However, for transcriptomic studies, scRNA-seq retains advantages in ease of application and scalability, and large numbers of scRNA-seq datasets have and will be generated on a range of biological samples. Therefore, the pairing of readily-applied multiplexed FISH technologies such as SABER with scRNA-seq will continue to serve as an important technical approach for the characterization of cellular gene expression states with spatial resolution.

Detection of complex pools of reporters or barcodes is another area where effective multiplexed FISH technologies can be applied. In cell culture, FISH-based barcode detection has been employed for the analysis of lineage,^62^ and barcode reading is an element of STARmap probe design.^39^ Here we demonstrate the utility of SABER for assaying the activity of isolated candidate enhancer sequences introduced *in vivo*. Simultaneous detection of reporter expression and cell type markers with single cell resolution is required for assessing cell-type specificity of CRM activity. Highly multiplexed reporter assays such as MPRAs^63, 63^ using bulk RNA-seq are useful for identification of active enhancers among complex introduced reporter libraries, but may combine multiple cell populations when applied to tissues with high cellular heterogeneity. SABER can be used to assign candidate active enhancers identified by such methods to specific marker-defined cell populations. Using SABER we were able to detect reporters across a broad range of expression levels, and to assay DNA plasmid copy number in the same cells, providing a tool to quantify enhancer strength and specificity. As an effective and simple method to robustly detect RNA and DNA sequences in cells and tissue, SABER enables the characterization of abundances, identities, and localizations of complex sets of endogenous and introduced nucleic acids.

## METHODS

### CONTACT FOR REAGENT AND RESOURCE SHARING

Further information and requests for resources and reagents should be directed to and will be fulfilled by the Lead Contact, Peng Yin.

### EXPERIMENTAL MODEL AND SUBJECT DETAILS

#### Cell culture

MRC-5 (human, ATCC CCL-171) were grown in Dulbecco’s modified Eagle medium (Gibco #10564) supplemented with 10% (vol/vol) serum (Gibco #10437), 50 U/mL penicillin, and 50µg/mL streptomycin (Gibco #15070). EY.T4 embryonic fibroblasts (mouse)^65^ were grown in Dulbecco’s modified Eagle medium supplemented with 15% (vol/vol) serum, 50 U/mL penicillin, and 50µg/mL streptomycin. All cells were cultured at 37°C in the presence of 5% CO_2_.

#### Tissue

All animal experiments were approved by the Institutional Care and Use Committees (IACUC) at Harvard University. Experiments were performed on tissue collected from wild-type male and female CD1 IGS mice (Charles River). Tissue were collected at postnatal day P17 for all experiments with the exception of electroporated retinas used for reporter assays (P13).

### METHOD DETAILS

#### DNA and RNA FISH probe design

Oligopaint FISH probe sets^44^ targeting mouse mRNAs, human chromosome 1, and the human X chromosome were discovered using the OligoMiner pipeline^49^ run with the ‘balance’ settings and accessed from the mm10 and hg38, respectively, whole-genome probe sets hosted at https://yin.hms.harvard.edu/oligoMiner/list.html. For RNA FISH probes, the genomic locations of the exons and/or introns of the target gene were acquired from the UCSC Genome Browser^66^ and used in combination with the ‘intersectBed’ utility of BedTools^67^ to isolate probe oligos targeting the RNA features of interest. As the aforementioned database of probe sequences exclusively contain ‘+’ strand information, in cases where the desired RNA target also carried ‘+’ strand sequence, the OligoMiner script ‘probeRC’ was used to convert the probe sequences to their reverse complements. Oligopaint probes targeting the six non-endogenous reporter RNA sequences and the reporter plasmid DNA backbone were also designed using OligoMiner, with the ‘blockParse’ script being run with ‘balance’ settings (-t 37 -T 42 -l 36 -L 41), alignment to the mm10 reference genome using Bowtie2^68^ with ‘–very-sensitive-local’ settings, then processed by ‘outputClean’ using ‘zero mode / −0’, and finally compared against a dictionary of all 18mers occurring in mm10 produced by Jellyfish 2.0,^69^ with candidate probes containing an 18mer occurring in mm10 >2 times being filtered. Oligo probes targeting the mouse major satellite, minor satellite, and telomere repeats were adapted from ref.^70^

#### 42mer bridge sequence design

The 42mer bridge sequences were drawn from blocks of orthogonal barcode sequences^50^ catenated together. They were then checked against target genomes using the same vetting process (Bowtie,^68^ Jellyfish^69^) used for candidate probe sequences (see above) as well as with BLAST.^71^ NUPACK^46–48^ (*ppairs* and *complexes* executable) were used to evaluate and subsequently screen the single-strandedness of probes and duplexing probabilities, respectively. See Supplemental Section 8 for a complete list of the designed 42mer bridge sequences.

#### PER primer sequence design

A set of 3-letter primer sequences were designed to have a minimum level of single-strandedness and a maximum probability of binding to existing 50mer concatemer sequences in the set, using NUPACK^46–48^) to calculate these prob-abilities and a custom Python optimization script. PER concatemeric sequences were then subjected to similar screening (with Bowtie,^68^ Jellyfish,^69^ and BLAST^71^) as the 42mer bridge sequences (see above). See Supplemental Section 8 for a complete list of probe-primer sequences and hairpins used.

#### DNA synthesis and purification

Probes and bridge sequences were typically ordered unpurified (standard de-salting) from IDT. The fluor-labeled primer used in Fig. 1b, fluorescent imager oligos, and many of the hairpins were HPLC purified. However, most primers can be successfully extended using unpurified hairpins bearing a 3’ poly-T sequence, and probes for tissue FISH were generated in this manner. Oligos were pre-suspended in 1×TE buffer (10mM Tris, 0.1 mM EDTA) at 100 µM or 200 µM concentration and diluted in 1 x×TE to a working concentrations of 10 µM. Stock and working solutions were stored at −20°C. Complex oligo libraries for Fig. 4b and Fig. 6 were ordered from Twist Bioscience (San Francisco, CA) and prepared as previously described.^33,44,72^ Briefly, libraries were first amplified using emulsion PCR, followed by a large-scale PCR and then *in vitro* transcription^33^ to generate RNA complements of probes. Reverse transcription and subsequent digestion of RNA produced single-stranded DNA probe sets.

#### PER concatemerization

Typically, 100 µL reactions were prepared with final concentrations of:1×PBS, 10mM MgSO_4_, 400 - 1000 units/ml of Bst LF polymerase (NEB M0275L or McLab BPL-300), 300 - 600 µM each of dATP/dCTP/dTTP (NEB M0275L), 100 nM of Clean.G hairpin, 50 nM - 1µM hairpin(s), and water to 90 µL (see Supplementary Fig. S1B for a visual representation). After incubation for 15 minutes at 37°C, 10 µL of 10 µM primer was added, and the reaction was incubated another 1-3 hours followed by 20 minutes at 80°C to heat inactivate the polymerase. PER extension solutions were directly diluted into ISH solutions or, in the case of combinatorial probe labeling, purified and concentrated using a MinElute (Qiagen #28004) kit or DNA Clean and Concentrator kit (Zymo, DCC-100) with distilled water elution to reduce volume and salt concentration from the reaction condition. Full extension and purification conditions and PER sequences for each experiment can be found in Supplementary Experimental Procedures.

#### Gel electrophoresis

Lengths of concatemers were evaluated by diluting 2 µL of *in vitro* reaction with 18 µL water and heat shocking at 95°C for 2 minutes. Samples were then run on 1% E-Gel EX agarose gels (Thermo Fisher G402001) for 10 minutes alongside a 1kB Plus Ladder (Invitrogen) and imaged with the Sybr Gold channel on a Typhoon FLA 9000 scanner.

#### Cell fixation

8 well chambers (Ibidi #80827) were seeded with MRC-5 cells and allowed to grow to desired confluency in a tissue culture incubator (37°C with 5% CO2). All subsequent steps were performed at room temperature, except where otherwise specified. After rinsing in 1×PBS, cells were fixed in a 1×PBS + 4% (wt/vol) paraformaldehyde solution for 10 minutes and then rinsed with 1×PBS. Chambers were then optionally stored at 4°C for up to two weeks before continuing with the ISH protocol.

#### DNA FISH in fixed cells

3D DNA FISH^73, 73^ closely followed previous protocols (see refs^18,44,49,72,75^). After a 1×PBS rinse (1 minute), samples were permeabilized for 10 minutes in 1×PBS with 0.5% (vol/vol) Triton X-100 and then for 2 minutes in 1×PBS + 0.1% (vol/vol) Tween-20 (PBT). After 5 minutes incubation in 0.1N HCl, samples were washed twice in 2×SSC + 0.1% (vol/vol) Tween-20 (SSCT) for (1×1 min, 1×2 min). After a further 5 minutes in 2×SSCT + 50% (vol/vol) formamide, samples were transferred to fresh SSCT + 50% (vol/vol) formamide and allowed to incubate at 60°C for at least one hour. Wells were subsequently loaded with 125 µL of ISH solution comprising 2×SSCT, 50% (vol/vol) formamide, 10% (wt/vol) dextran sulfate, 40 ng/ µL RNase A (EN0531, Thermo Fisher), and each PER extension at ∼67 nM final concentration (1:15 dilution from 1µM PER) and denatured at 80°C for 3 minutes. After overnight incubation at 44°C on a flat-block thermocycler (Eppendorf Mastercycler Nexus), 200 µL pre-warmed 2×SSCT (at 60°C) was added, and the hybridization solution was aspirated. Samples were washed for a further 20 minutes (4×5 minutes) in 2×SSCT at 60°C and 4 minutes (2×2 minutes) in 2×SSCT at RT. Samples were then transferred to 1×PBS and washed a couple times (2×2 minutes and then fresh solution) before continuation with the fluorescent hybridization protocol.

#### RNA FISH in fixed cells

RNA FISH was performed similarly to 3D DNA FISH, but with a shortened protocol. After a 1×PBS rinse (1 minute), samples were permeabilized for 10 minutes in 1×PBS with 0.5% (vol/vol) Triton X-100 and then for 1 minutes in 1×PBS + 0.1% (vol/vol) Tween-20 (PBT). Samples were then transferred to 2×SSCT (1 minute) before wells were loaded with 125 µL of ISH solution comprising 2×SSCT, 50% (vol/vol) formamide, 10% (wt/vol) dextran sulfate, and PER extension at ∼67 nM final concentration (1:15 dilution from 1µM PER). After a 3 minute denaturation at 60°C, chambers were incubated overnight 42°C on a flat-block thermocycler (Mastercycler Nexus, Eppendorf). After hybridization, 200 µL pre-warmed 2×SSCT (at 60°C) was added, and the hybridization solution was aspirated. Samples were washed for a further 20 minutes (4×5 minutes) in 2×SSCT at 60°C and 4 minutes (2×2 minutes) in 2×SSCT at RT. Samples going directly to fluorescent hybridization were then transferred to 1×PBS and washed (1 minute and then fresh solution). Samples were optionally held at 4°C (1-2 overnights) before continuation with branching or fluorescent hybridization protocols.

#### Metaphase DNA FISH

Human metaphase chromosome spreads on slides (XX 46N or XY 46N, Applied Genetics Laboratories) were denatured in 2×SSCT + 70% (vol/vol) formamide at 70°C for 90 seconds before being immediately transferred to ice-cold 70% (vol/vol) ethanol for 5 minutes. Samples were then immersed in ice-cold 90% (vol/vol) ethanol for 5 minutes and then transferred to ice-cold 100% ethanol for a further 5 minutes. Slides were then air-dried before 25 µL of ISH solution comprising 2×SSCT, 50% (vol/vol) formamide, 10% (wt/vol) dextran sulfate, 40 ng/ µL RNase A (EN0531, Thermo Fisher), probe pools with bridges at ∼500 nM, and PER extension at 96 or 192 nM final concentration was added. Rubber cement was used to seal the hybridization solution underneath a coverslip, and the sample was placed into a humidified chamber inside an air incubator at 45°C overnight. After hybridization, samples were washed in 2×SSCT at 60°C for 15 minutes and then in 2×SSCT at room temperature (2×5 minutes). See Supplemental Experimental Procedures for additional information.

#### Branch hybridization

After washing the samples in 2×SSCT at room temperature (2 minutes), branch hybridization solutions comprising 2×SSCT, 30% (vol/vol) formamide, 10% (wt/vol) dextran sulfate, and PER extension at ∼67 nM final concentration (1:15 dilution from 1µM PER) were added. After hybridization at 37°C for 1.5 hours, 200 µL pre-warmed 2×SSCT (at 60°C) was added, and the hybridization solution was aspirated. Samples were washed for a further 20 minutes (4×5 minutes) in 2×SSCT at 60°C and 4 minutes (2×2 minutes) in 2×SSCT at room temperature. Samples were then transferred to 1×PBS and washed (1 minute and then fresh solution). For iterative branching experiments, slightly less stringent conditions were used: 30°C for 1 hour (instead of 37°C for 1.5 hours) for the hybridization, 55°C (instead of 60°C) for the heated washes, and only one of the two 2×SSCT washes at room temperature. This hybridization and washing was performed repeatedly, with wells not receiving hybridization solution held in 2×SSCT. After branching steps, samples transferred to 1×PBS and washed (1 minute and then fresh solution). Samples were then typically held at 4°C overnight before continuation with fluorescent hybridization protocols.

#### Fluorescent hybridization (chamber)

After washing once in 1×PBS, 125 µL fluorescent hybridization solution comprising 1×PBS and 1 µM fluorescent imager strands was introduced to each well. After incubation for 1 hour at 37°C, samples were washed in pre-warmed 1×PBS 3 times (5 minutes and 2×2 minutes) at 37°C. After a final rinse with 1×PBS at room temperature 1×PBS, SlowFade Gold + DAPI (Thermo Fisher S36939) was added for diffraction-limited imaging.

#### Fluorescent hybridization (metaphase slide)

For spectral imaging (Fig. 4b), slides were transferred to 1×PBS and then dried before 25 µL fluorescent hybridization solution comprising 1×PBS with 1 µM fluorescent imager strands was added. After covering the hybridization solution with a coverslip, slides were put into a humidified chamber and incubated in an air incubator at 37°C for 1 hour. Slides were then washed three times (1×15 minutes and 2×5 minutes) in pre-warmed 1×PBS at 37°C before being dried. 12 µL of SlowFade Gold + DAPI (Thermo Fisher S36939) was added and sealed underneath a coverslip with nail polish before imaging.

#### Fluorescent exchange (metaphase slide)

For the metaphase walk (Fig. 6), slides were transferred to 1×PBS and then dried. A flow chamber (with volume ∼50 µL) was then constructed using a coverglass attached to the slide with double-sided tape to allow fluid exchange, and samples were re-hydrated in 1×PBS. Each hybridization comprising 1×PBS with 10% (wt/vol) dextran sulfate and 1 µM fluorescent imager strands was incubated at room temperature for 15 minutes. (For combinatorial hybridizations with PER concatemers mapped to two colors, each of those fluorescent imager strands was included at 500 nM each, retaining the 1 µM overall concentration). After washing with 200 µL (∼4×flow through the chamber), SlowFade Gold + DAPI (Thermo Fisher S36939) was added for diffraction-limited imaging. Between each fluorescent hybridization, previous imager strands were stripped with formamide as follows. First, SlowFade was washed out with 200 µL of 1×PBS, followed by stripping with a total of 1.6ml of 1×PBS containing 60% (vol/vol) formamide flowed through over the course of 15 minutes. After stripping, slides were washed with 200 µL 1×PBS, allowed to sit for 2 minutes, then washed twice more with a total of 400 µL 1×PBS before adding the next fluorescent hybridization solution. The slide was stored overnight at 4°C overnight after the first six hybridizations before subsequent stripping and hybridization steps.

#### Retinal histology

Neural retinas were dissected in PBS and fixed for 25 minutes at room temperature in 4% formaldehyde solution (diluted in 1×PBS from 16% methanol-free formaldehyde solution (ThermoScientific 28908)). For cryosectioning, retinas were transferred to a solution composed of 50% O.C.T. and 15% sucrose in 0.5×PBS and frozen in an ethanol bath prior to long-term storage at −80°C. Cryosections cut to 35 or 40 µm were adhered to Poly-D-Lysine-coated (Sigma P6407) 8-well Ibidi chamber slides and dried. Prior to hybridization tissue were washed in PBS with Tween-20 (Sigma P9416) at.1% (vol/vol), then received pretreatment consisting of a mild proteinase K exposure (1.5 µg / ml, 15 minutes) followed by post-fixation and acetic anhydride treatment as described previously.^53^ Sections were incubated at 43°C in a hyb oven in wash hyb (40% formamide, 2×SCC pH 7, 1% Tween-20) for 30 minutes preceding addition of pre-warmed probe/hyb solution. Probe concentrations were determined by nanodrop and probes were added to a final mass of 1µg each per well (120 µL volume) in hyb solution (40% formamide, 2×SSC pH 7, 1% Tween-20, and 10% dextran sulfate (Sigma, D8906)). Following overnight incubation (18-24 hours), slides were washed 2×30 minute in 40% formamide wash hyb, 2×45 minutes in 25% formamide wash hyb, and 2×15 minutes in 2×SSCTw (0.1% tween). For branching, the 27*.27*.27*.28 branch was extended to a length of 500 nt and incubated for at least 5 hours in hyb solution at 37°C. Washes were performed as for the primary probe incubation with temperature set to 37°C. For fluorescent detection slides were washed three times in PBSTw at room temperature, and then transferred to 37°C for hybridization and subsequent wash steps. Detection oligos were diluted to a concentration of 1 µM in a 1×PBS solution with 0.2% Tween-20 and 10% dextran sulfate. This solution was incubated with the sample for 2 hours, and then washed 4×7 minutes in PBSTw. Imaging was performed in 80% glycerol mounting media (80% glycerol, 1×PBS, 20mM Tris pH 8, and 2.5mg/mL of propyl gallate). For serial detections, fluorescent oligos were stripped with a solution of 50% formamide in 1×PBS at room temperature (3×5 minutes washes), and washed 3×2 minutes in PBSTw. For DNAse digestion of primary probe, after 1st detection and formamide stripping of imager oligos, samples were washed 3×5 minutes in PBSTw, once in DNAse I buffer (Sigma, 04716728001), and then incubated in 40U of DNAse I (Sigma, 04716728001) or 80U Exonuclease I (NEB, M0293S), representing 1:50 dilution of each enzyme in DNAseI buffer, with 200ul total volume. Samples were incubated 30 minutes as 37°C and then washed at room temperature 3 times in PBS with 5mM EDTA, fixed 10 minutes in 4% FA in PBS, and washed 3 times in PBSTw before proceeding with the second overnight probe hybridization.

#### Whole mount retinal staining

Whole mount stainings were conducted with a similar protocol but with extended hybridization and wash times. Primary probe was incubated for 32 hours, followed by 2×45 minutes wash in 40% formamide wash hyb, 2×90 minutes wash in 25% formamide wash hyb, and 2×20 minutes in 2×SSCTw. 200 µL volumes were used for hybs and washes, and 1.8 µg of *Grik1* probe (500 nt length) was used. Fluorescent oligos were incubated for 8 hours followed by 3×30 minutes wash in PBSTw (0.2% Tween). Retinas were flattened by creating 4 incisions prior to fixation, and underwent pretreatment as described above while floating. Next, flattened retinas were transferred to an Ibidi chamber slide with ganglion cell layer facing up. Retinas were held in place by application of a nylon mesh (SEFAR-NITEX 03-64/45) cut to size and layed over the retina. The corners of the mesh were glued to the corners of the well using White Gorilla Glue. Prior to this step, retinas were transferred to O.C.T./sucrose solution (see above) to protect tissue from dessication while the mesh was overlayed and glue was dried. Prior to application of the first hyb solution, O.C.T. solution was washed away with 3×10 minute washes in PBS with.3% Triton-x (Sigma, T8787) followed by 2×30 seconds washes in ddH20.

#### Retina DNA FISH

For detection of DNA, retinas were treated as described for RNA FISH with the addition of a 5 minute treatment in a 1N HCl, 0.5M NaCl solution, followed by a 15 minute incubation at 80°C in a solution of 50% formamide and 2×SSC on a preheated metal block prior to the primary probe hyb. For RNA and DNA co-detection, DNA denaturation and hybridization was conducted following RNA detection. RNAse A (Thermo Fisher EN0531) was added to the primary probe hybridization solution for the DNA FISH step at a concentration of 200 ng/µl.

#### Retina immunohistochemistry and WGA counterstain

WGA conjugated to 405s (Biotium, 29027) was diluted to a concentration of 10 µg/mL in PBSTw and samples were incubated for 1 hour following each round of fluorescent oligo detection. Slides were washed 2 × 5 minutes in PBSTw following WGA application. Antibodies were applied following FISH detection. Slides were pre-incubated in block (5% HIDS, 0.3% Triton-X in PBS) for 1 hour. Anti-PKCa at 1:1500 (Sigma, P4334) and anti-Calretinin at 1:1000 (Abcam, AB1550) were incubated overnight at 4°C in block, washed 4 times in PBSTx (PBS with 0.3% Triton-X) over the course of 2 hours, incubated for at least 4 hours in secondary antibody (1:500), and then washed for 30 minutes in PBSTx prior to addition of mounting media and imaging. For GFP reporters, GFP signal was amplified with chicken anti-GFP (Abcam, AB13970) at 1:1000. Secondary antibodies used were: Donkey anti-Goat Alexa 647 (Jackson Imm-munoRes 705-605-147), Donkey anti-Chicken Alexa 488 (703-545-155), and Donkey anti-Rabbit Alexa 488 (711-545-152).

#### Reporter construct cloning and electroporation

Reporter sequences were derived from full or partial sequences of genes commonly expressed heterologously in mammals: dCas9 (template: Addgene #60954), Lacz, Cre, and Luciferase (see Supplemental Experimental Procedures for full list of sequences). Reporter mRNAs were designed to be within a size range of 1-1.6kb and targetable by 22-24 primary probes. We used a reporter plasmid with a minimal TATA promoter (Stagia3),^57^ replacing the GFPiAP ORF of this plasmid with the described reporter sequences by plasmid digestion with Age1/EcoRV followed by insertion of reporter sequences in frame using Gibson assembly and transformation into DH5alpha cells. For reporter validation, a Vsx2 (Chx10) enhancer^57^ was inserted upstream of the TATA box in each re-porter plasmid at the EcoRI site using gibson assembly. For reporter screening, each of the six regions corresponding to open chromatin peaks was amplified using PCR with purified Mus musculus genomic DNA as template. PCR products were inserted upstream at the EcoRI site of reporter plasmids. DNAse I hypersensitive regions for 1 week adult retina^58^ were identified and displayed using the UW DNAse I HS track in the UCSC genome browser. Reporters were electroporated into mouse pups via subretinal injection at P1 as described^59^ at a concentration of 500 ng/µl for each construct. For experiments with DNA FISH detection of plasmid, the CRM-4 reporter plasmid was electroporated at a concentration of 1.5 µg/ul. CAG-nls-tagBFP or CAG-mtagBFP (membrane localized) were co-electroporated at a concentration of 200 ng/µl to enable identifi-cation and orientation of electroporated regions prior to sectioning. For plasmid DNA FISH, pENTR/pSM2(CMV) GFP (Addgene #19170) plasmid was used as the co-electroporation marker at a concentration of 100 ng/µl. This plasmid has little sequence similarity to the Stagia3 backbone and is minimally targeted by the probe set used for detection of Stagia3. For validation of CRM-1 and CRM-4 using a protein-based (GFP) reporter assay these enhancer sequences were re-cloned into Stagia3.

#### Microscopy

Imaging of iterative branching samples was conducted on an inverted Zeiss Axio Observer Z1 using a 100x Plan-Apochromat Oil N.A. 1.40 objective. Samples were illuminated by using Colibri light source using a 365 nm, 470 nm, 555 nm, or 625 nm LED. A filter set composed of a 365 nm clean-up filter (Zeiss G 365), a 395-nm long-pass dichroic mirror (Zeiss FT 395), and a 445/50 nm band-pass emission filter (Zeiss BP 445/50) was used to visualize DAPI staining. A filter set composed of a 470/40 nm excitation filter (Zeiss BP 470/40), a 495 nm long-pass dichroic mirror (Zeiss FT 495), and a 525/50-nm band-pass emission filter (Zeiss BP 525/50) was used to visualize ATTO 488 signal. A filter set composed of a 545/25-nm excitation filter (Zeiss BP 545/25), a 570 nm long-pass dichroic mirror (Zeiss FT 570), and a 605/70 nm band-pass emission filter (Zeiss BP 605/70) was used to visualize ATTO 565 signal. Finally, a filter set composed of a 640/30-nm excitation filter (Zeiss BP 640/30), a 660 nm long-pass dichroic mirror (Zeiss FT 660), and a 690/50 nm band-pass emission filter (Zeiss BP 690/50) was used to visualize Alexa Fluor 647 signal. Images were acquired by using a Hamamatsu Orca-Flash 4.0 v3 sCMOS camera with 6.5 µm pixels, resulting in an effective magnified pixel size of 65 nm. Images were processed by using Zeiss ZEN software and Fiji/ImageJ.^54, 54^

Remaining cell and metaphase samples imaged on a Nikon Eclipse Ti-E microscope by using a CFI PlanApo 100x Oil (N.A. 1.45) objective. Illumination was performed with a Spectra X LED system (Lumencor) using a 395/25 nm, 295 mW LED for DAPI signal, a 470/24 nm, 196 mW LED for ATTO 488 signal, a 550/15 nm, 260 mW LED for ATTO 565 signal, and a 640/30 nm, 231 mW LED for Alexa Fluor 647 signal. Illumination light was spectrally filtered and directed to the objective, and emission light was spectrally filtered and directed to the camera by one of four filter cubes: 1) Semrock BFP-A-Basic-NTE for DAPI, 2) Semrock FITC-2024BNTE-ZERO for ATTO 488, 3) Semrock TRITC-B-NTE-0 for ATTO 565, and 4) Semrock Cy5-4040C-NTE-ZERO for Alexa 647 signal. An Andor Zyla 4.2+ sCMOS camera was used to acquire images with 6.5 µm pixels, resulting in an effective magnified pixel size of 65 nm.

All tissue images were acquired on a Zeiss Axio Observer Z1 inverted microscope equipped with an LSM780 single point scanning confocal attachment that contains two Quasar alkali PMTs and a GAaSP 32 channel spectral detector. Images were acquired using either a Plan Apo 40x/1.3 DIC or Plan Apo 63x/1.4 DIC oil objective. Laser lines used were 405, 488, 561, 594, and 633. Dichroic and main bean splitters used were MBS458, MBS488, MBS488/561/633, 405. The imaging software was ZEN Black 2012.

#### Image processing

Maximum projections taken on the Nikon Eclipse Ti-E microscope were processed using the Nikon Elements software, and by Zeiss ZEN software for non-confocal images taken on the Zeiss Axio Observer 1. Images were then processed with Fiji/ImageJ.^76^ Multicolor overlays of cells and metaphase spreads were generated using a Python script written to mimic the ‘screen’ behavior of Photoshop, which also allowed automatic cropping, contrasting, and DAPI alignment. Most images presented in the main and supplemental figures utilized max projections of Z-stacks, with the exception of the metaphase spread image in Fig. 4b, the interphase image in Fig. 4b, and the metaphase spreads in Fig. 6, for which single in-focus z slices for each hybridization were utilized to create overlays. For cell images with nuclei outlined, nuclear outlines were first automatically generated using CellProfiler^51, 51^ analysis pipelines (see below), and then these outlines were automatically outlined and then re-styled in Adobe Illustrator. Scale bars were added either in Adobe InDesign or Adobe Illustrator based on expected pixel size scaling. For retina images, maximum intensity projections were generated in ZEN 2.3 lite. Multicolor overlays were generated using the screen setting in Adobe Photoshop, and brightness and contrast were adjusted for display purposes using Adobe Photoshop or Fiji/ImageJ.^76^ For whole mount volume visualizations the ImageJ plugins 3D Viewer and Volume Viewer were employed. For quantification of intensities and puncta detection in retina tissue, MATLAB and the Image Processing Toolbox were used (MATLAB and Image Processing Toolbox Release R2018a, The MathWorks, Inc., Natick, Massachusetts, United States). Open Microscopy Environment’s Bio-Formats library^78^ was used for image file manipulation, including import of image stacks to the MATLAB environment.

For experiments with serial detection and imaging, images from the same retinal regions were aligned based on the WGA stain using the intensity-based automatic image registration tools in the MATLAB Image Processing Toolbox.

To subtract tissue autofluorescence for the 9×retina overlay, puncta were first detected in 3D by the described method (see below) and a Gaussian filter slightly larger than that used for puncta detection was convolved with the image of puncta centroids to capture all voxels in the puncta. The resulting mask was applied to the original SABER image, yielding a background-subtracted version of the original image while preserving the original signal pattern.

For high-resolution renderings of detected puncta (as shown in Fig. 7J), images of detected puncta centers were resized to a resolution of *∼*10nm/pixel using bicubic interpolation and then spherically dilated to size similar to the original puncta, which can be estimated using the *Analyze* → *Measure* functions in ImageJ based on the original SABER images.

In all cases except where otherwise noted in figure legends, images were only contrasted to improve signal visibility by changing the min (black) and max (white) values.

### QUANTIFICATION AND STATISTICAL ANALYSIS

#### Puncta quantification in cells

Maximum intensity projections in Z were created using Nikon Elements soft-ware from raw mutlichannel Z-stacks. These max projections were then inputted into CellProfiler 3.0,^51, 51^ in which an automated image analysis pipeline was constructed to identify nuclei, cell bodies, and FISH foci and to calculate background-subtracted maximum pixel intensity of each segmented focus. For intensity quantification experiments, the same pipeline was used for all conditions being compared. For cases where the number of puncta per cell or nucleus was calculated, a parent-child relationship was established between the FISH foci and the respective cellular or subcellular feature. For fold enhancement calculations in Fig. 2BD and Fig. S2H, background was calculated as the mean of the image pixels masked for the detected puncta. Background-subtracted peak intensity distributions of puncta were then divided by the average of the corresponding distribution for the unextended condition. Cumulative density distributions depicted in Fig. 2G did not subtract background for fold enhancement calculations because the sample was very crowded. In Fig. 4D, co-localization of puncta was assumed if any pixels within the detected area of a punctum in one channel overlapped with any pixels corresponding to a punctum detected in the other channel. See Supplemental Experimental Procedures for cell and puncta numbers.

#### Drop-seq data processing

Bipolar cell Drop-seq data^53^was processed according to the mark-down accompanying the manuscript using class file class.R (provided at https://github.com/broadinstitute/BipolarCell2016). All 10,888 cells identified present in cluster 1 (corresponding to Rod bipolar cells) following Louvain clustering and cluster merging were used for plotting average number of transcripts per cell. This analysis discards cells considered of poor quality with fewer than 500 detected genes per cell. Dot plots were generated using the dot.plot function with use of all validated retinal cell type clusters from this dataset, as described in the markdown.

#### 3D retinal cell segmentation and alignment

To segment cells in retinal tissue, we applied an open-source membrane-based segmentation software, *ACME*,^52^ using the WGA signal as a membrane stain. Segmented images were manually masked in ImageJ to remove the inner plexiform layer and photoreceptor inner segments. Size filters were applied to remove improperly segmented or non-cellular structures. For serially imaged retina regions, cell segmentation was performed on the WGA channel from a single development and was applied to all registered channels. After automated segmentation, results were verified and visualized using the open-source ITK-SNAP software.^79^

#### 3D puncta detection and cell calling

Proper assignment of fluorescent SABER puncta to cells in tissue required detection of puncta in 3D. SABER images were processed using custom MATLAB software for localization of fluorescent puncta using a Laplacian of Gaussian method,^80^ similar to the analagous 2D pipeline implemented in.^10^ Briefly, this involves convolving the original SABER image with a noise-suppressing Gaussian filter with filter size corresponding to the empirical size of SABER puncta. The filter is elliptical in the Z-dimension to account for the point spread function. The Laplacian of the Gaussian-filtered SABER image was then taken to enhance signal detection and a threshold was set to differentiate from remaining background in a semi-automated way (see Fig. S3B-C and Data and Software Availability section for details).

For intensity quantification experiments in tissue, puncta centers were detected as described and were dilated to encompass the original puncta pixels. The maximum pixel intensity of each puncta was then taken. Average background pixel intensity was calculated by taking the average pixel intensity of the image masked by the complement of a spherically dilated image of detected puncta centroids (radius = 2 µm). To generate background-subtracted intensity distributions for fold enrichment calculations (Fig. 3C, L), the average background intensity was subtracted from the maximum intensity value of each puncta in the image. See Supplemental Experimental Procedures for puncta and section numbers.

A universal threshold was applied to call cells positive for each marker based on the distribution of puncta per cell for that transcript across all cells (see Fig. S3D for details). Similarly, for the bipolar probes, thresholds were 15, 5, and 2 for *Prkca, Tpbg*, and *Slc4a*, respectively. For quantification of re-porter RNA versus plasmid DNA, a threshold of 2 puncta per cell was used for CRM-4 reporter RNA, 7 puncta per cell for endogenous *Grik1*, and 3 puncta per cell for plasmid DNA. This method of analysis demonstrates quantification of puncta per cell using single markers and a universal cell outline, which can be useful in cases where cell-specific outlines and combinatorial marker sets are not yet available. The multiplexing capability of SABER (see Fig. 4) permits use of multiple markers for intersectional cell classification, which can alleviate inevitable thresholding ambiguity that arises due to both true biological variance and to segmentation imperfections.

#### Reporter specificity analysis

To quantify the specificity of each reporter, we used the DNA FISH image stacks to estimate that 52% of electroporated cells were rods, 9% were positive for Grik1 endogenous RNA, 18% were ON bipolar cells, 12% M¨uller Glia, and 9% amacrine cells. These numbers were estimated directly from the data based on plasmid DNA detection, Grik1 RNA expression, and the known cell body localization of each cell type. To calculate reporter specificity and statistical significance, we used a hypergeometric test^81^ to evaluate the probability of observing the empirical positive cell patterns for each CRM. For a given CRM-driven reporter (CRM 1-6) and endogenous gene (Rho, Grm6, Grik1, Glyt1/Gad1, Other), we can consider *C*_*G*+_, the number of CRM reporter-positive cells that are positive for the endogenous gene:

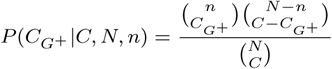

Where,

CRM^+^ = Set of cells positive for CRM-driven reporter RNA

GENE^+^ = Set of cells positive for endogenous gene

*C_G_*+ = |CRM^+^ ∩ GENE^+^| = Total # cells positive for both the CRM and gene

*N* = Total # cells that received CRM-reporter plasmid DNA

*n* = Total # GENE^+^ in the plasmid-receiving population

*C* = |CRM^+^| = Total # cells observed positive for CRM reporter RNA

We took *N* =1,500 as an estimate of the total cell population assayed for each CRM based on the number of plasmid-positive cells observed in the DNA FISH experiment. In any one 240 µm×240 µm electroporated retinal region, we estimate that approximately 300 cells received plasmid DNA based on the automated cell segmentation. Cells were analyzed across 5 similar retinal regions, yielding a total population size of approximately 1,500 cells.

With this estimate, *n* can be inferred based on the proportion of each endogenous marker within the electroporated population. Both *C* and *C*_*G*+_ were directly measured.

#### Plotting and visualization

Most plots and some image overlays were generated in Python, using the Matplotlib,^82^ Seaborn,^83^ NumPy,^84^ Pandas,^85^ PIL, and Biopython^86^ libraries. Data was imported either in CSV format or read in from CellProfiler^51, 51^ output files. The plot in Fig 7I was generated using the *ggballoonplot* function of ggpubr,^87^ a package for ggplot2^88^ in R.^89^

### DATA AND SOFTWARE AVAILABILITY

The complete set of CellProfiler^51, 51^ pipelines used as well as example input images for each are available at https://github.com/brianbeliveau/SABER. *PD3D*, a package of MATLAB functions for detecting SABER puncta (or other fluorescent puncta) in 3D and assigning puncta to cells in a watershed segmentation is available at https://github.com/ewest11/PD3D. Any remaining scripts for image processing and plotting will be made available upon request. Step-by-step protocols will be posted online at http://saber.fish or http://saber-fish.net/.

## Acknowledgments

The authors thank Brandon Fields, Scott Kennedy, Jumana Alhaj Abed, Ting Wu, Tom Ferrante, Ninning Liu, Frits Dannenberg, Marcelo Cicconet, and Paula Montero Llopis for fruitful discussions and technical support. This work was supported by the Office of Naval Research (under grants N00014-13-1-0593, N00014-14-1-0610, N00014-16-1-2182, and N00014-16-1-2410), the National Institutes of Health (under grants 1R21HD072481-01, 1R01EB018659-01, 1-U01-MH106011-01, 5K99EY028215-02), the National Science Foundation (under grant CCF-1317291), the Howard Hughes Medical Institute, and the Wyss Institute’s Molecular Robotics Initiative (MRI). J.Y.K. was supported by a National Science Foundation Graduate Research Fellowship, B.J.B. was supported by a Damon Runyon Cancer Research Foundation Fellowship (HHMI), S.W.L. was suported by HHMI and the National Institutes of Health (grant 5K99EY028215-02), E.W. was supported by a Genetics and Genomics PhD Training Grant (NIH T32 GM096911), H.M.S. was supported by a Uehara Memorial Foundation Research Fellowship, and S.K.S. was supported by a long-term postdoctoral fellowship from Human Frontier Science Program (HFSP) (LT000048/2016-L) and an EMBO long-term fellowship (ALTF 1278-2015).

## Author Contributions

Conceptualization, J.Y.K., B.J.B., S.W.L., C.L.C., and P.Y.; Methodology, J.Y.K., B.J.B., S.W.L., and E.W.; Software, J.Y.K., B.J.B., and E.W.; Validation, J.Y.K., B.J.B., S.W.L., A.Z., and S.K.S.; Formal Analysis, J.Y.K., B.J.B., S.W.L., and E.W.; Investigation, J.Y.K., B.J.B., S.W.L., E.W., A.Z., H.M.S., S.K.S., and Y. W.; Resources, J.Y.K., B.J.B., S.W.L., H.M.S., and S.K.S.; Data Curation, J.Y.K., B.J.B., S.W.L., and E.W.; Writing – Original Draft, J.Y.K., B.J.B., S.W.L., and E.W.; Writing -Review and Editing, J.Y.K., B.J.B., S.W.L., E.W., C.L.C., and P.Y.; Visualization, J.Y.K., B.J.B., S.W.L., and E.W.; Supervision, C.L.C. and P.Y.; Project Administration, C.L.C., P.Y.; Funding Acquisition, C.L.C. and P.Y.

